# In the brain of the beholder: whole brain dynamics shape the perception during ambiguous motion

**DOI:** 10.1101/2025.02.05.636650

**Authors:** Alessandra Pizzuti, Irene Acero Pousa, Omer Faruk Gulban, Judith Peters, Gustavo Deco, Rainer Goebel

## Abstract

Visual perception is typically based on a one-to-one mapping between stimuli and conscious experiences. However, under bistable conditions, identical sensory inputs can elicit alternating perceptions, requiring the brain to resolve ambiguity. The mechanisms underlying transitions between distinct perceptual states and their sustained maintenance remain poorly understood. In this ultra-high-field (7T) fMRI study, we investigated the neural dynamics of perception using a bistable motion stimulus (ambiguous motion quartet) that evoked endogenous alternations between horizontal and vertical motion, compared to a control condition (physical motion quartet) with unambiguous sensory input. Consistent with previous findings, the human motion complex (hMT+) played a central role in processing both physical and ambiguous motion conditions. By dissociating neural activity during perceptual transitions from sustained perceptions, we found that hMT+ mostly interacts dynamically with area 46 in the frontal cortex and PF/PFm within the inferior parietal lobe during transitions and with subregions of the superior parietal lobe during sustained perceptions. Beyond local activity, computational modeling revealed an increase in hierarchical organization across cortical networks during the ambiguous condition. In particular, the same frontal and parietal regions exhibited ascension within the functional hierarchy, likely reflecting their specific role in coordinating computations for resolving ambiguity.

## 1 Introduction

Conscious perception is considered a hallmark of advanced mammals, reflecting the evolution of neural systems that bridge sensation and action. In humans, the progressive corticalization of these systems supports the ability to process ambiguous sensory inputs, enabling the brain to infer meaning and stabilize perception amidst uncertainty (Mesulam, 1998). This complexity is mirrored in the brain’s hierarchical organization, where networks of regions collaborate to optimize communication and computation (Baars, 1988; Fuster, 2022). Evidence suggests that different conditions can dynamically reconfigure the functional hierarchy of the brain to adapt to varying cognitive and computational demands (Deco et al., 2022, 2024; M. Kringelbach et al., 2022). An approach to studying the neural mechanisms underlying conscious perception is through bistable stimuli, which require the brain to resolve ambiguous inputs, resulting in endogenous perceptions that alternate between two competing interpretations (J. Brascamp et al., 2018; Leopold and Logothetis, 1999). Functional magnetic resonance imaging (fMRI) and transcranial magnetic stimulation (TMS) studies in humans have consistently reported the involvement of the fronto-parietal brain network in perceptual transitions during multistable tasks (J. Brascamp et al., 2018). This network is also associated with cognitive functions such as visual working memory (Marois and Todd, 2004), perceptual decision-making (Heekeren et al., 2004), spatial attention (Silver et al., 2005; Yantis et al., 2002), and guiding eye movements (Corbetta et al., 1998). More specifically, Brascamp and colleagues (J. Brascamp et al., 2018, Figure 3) highlight dorsolateral prefrontal cortex (DLPFC), frontal eye fields (FEF), temporoparietal junction (TPJ), intraparietal junction (IPJ), intraparietal sulcus (IPS), and inferior frontal cortex (IFC) as hotspots of consistent activations during bistable perception resulting from a meta-analysis across multiple fMRI and TMS studies. While there is broad consensus on the involvement of this network in bistable perception, the specific roles of individual regions and subregions in resolving bistable stimuli remain an ongoing debate. Specifically, it remains unclear whether the neural mechanisms that drive perceptual transitions between competing interpretations are the same as those that sustain a single interpretation over time. Furthermore, how these processes manifest within the brain’s hierarchical organization, particularly in response to the computational demands of ambiguity resolution, has yet to be fully explored. To address this, we conducted a 7T fMRI study using an ambiguous motion paradigm, where participants experienced alternating perceptions of horizontal and vertical motion under constant sensory input (**Figure 1**). Responses to this ambiguous condition were compared with a control condition using a physical motion stimulus that regularly induced alternating perceptions via unambiguous input. Additionally, resting-state fMRI data were collected to examine differences in the brain’s intrinsic functional architecture between rest and task conditions. Consistent with previous findings, we observed a critical role for the human motion complex (hMT+) in processing motion information across both conditions (Goebel et al., 1998; Muckli et al., 2002; Pizzuti et al., 2024; Salzman et al., 1992; Schneider et al., 2019; Sterzer et al., 2002). By dissociating neural responses during perceptual transitions from those during sustained perception in the ambiguous condition, we found that hMT+ dynamically interacts with area 46 in the frontal cortex and PF and PFm in the inferior parietal lobe during transitions and with subregions of the superior parietal lobe during sustained perception (**Figure 2**, **Figure 3**, **Figure 4**, **Figure 5**). All region names follow the Glasser parcellation (Glasser et al., 2016). Expanding beyond individual brain areas, we leveraged computational models to examine whole-brain hierarchical organization. Our analysis revealed that resolving ambiguity and constructing a coherent percept required increased hierarchical organization across the cortical network compared to both the physical motion condition and resting-state condition (**Figure 6**). Notably, the same frontal and parietal areas identified earlier were observed to ascend within the hierarchy during the ambiguous condition, reflecting their dynamic role in coordinating computations within the network necessary for perceptual resolution (**Figure 7**). This finding underscores the brain’s ability to adaptively reconfigure its hierarchical organization to meet complex computational demands.

**Figure 1:**
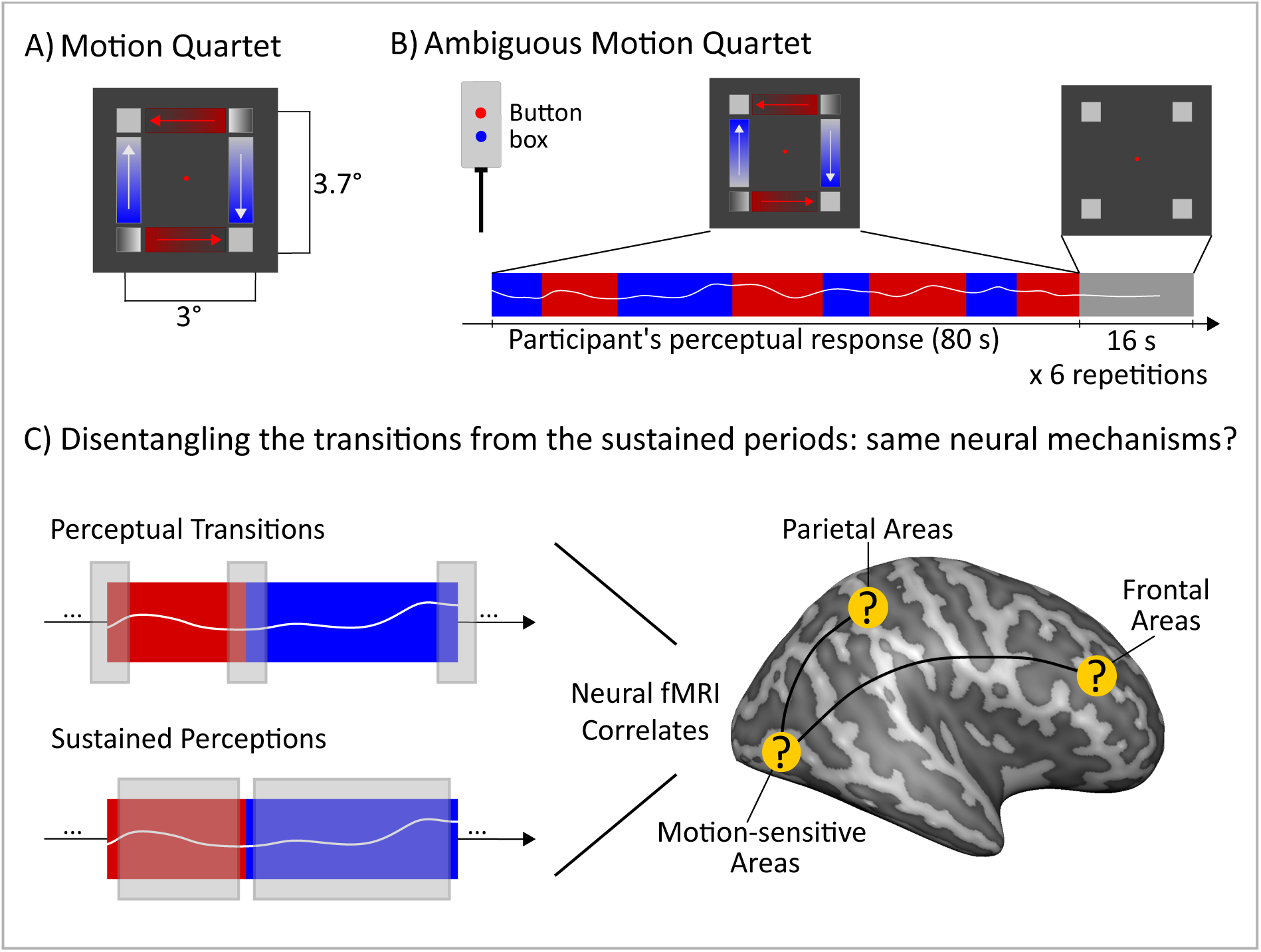
Stimulus, paradigm and experimental question for the ambiguous motion condition. A) Motion Quartet stimulus. Distances between inducers are in visual degrees. The two coupled inducers along the two diagonals alternate at 2.3 Hz. The gradient of gray filling two inducers indicates the pair in this schematic while the assigned color is gray (Animations: https://doi.org/10.6084/m9.figshare.21908394). B) Schematic illustration for an ambiguous motion run: while retinal input remains constant, the elicited perception of motion direction endogenously alternates between horizontal (red) and vertical (blue). Participants indicate the moment when perceptual transitions occur using an MRI-compatible button box. C) Disentangling the brain correlates of transition perception (switch) from sustained perception periods (block). Example of snapshot of a time course for the ambiguous motion condition where an horizontal motion perception (red) is followed by a vertical motion perception (blue). Gray squares indicate the two phases of the perception, respectively, the perceptual transitions (top row) and sustained perception (bottom row). Although the motion-sensitive areas as well as areas from the fronto-parietal network are expected to be involved when processing bistable stimuli, the specific functional roles of these areas during perceptual transitions and sustained perceptions and their interactions are still unknown.

**Figure 2:**
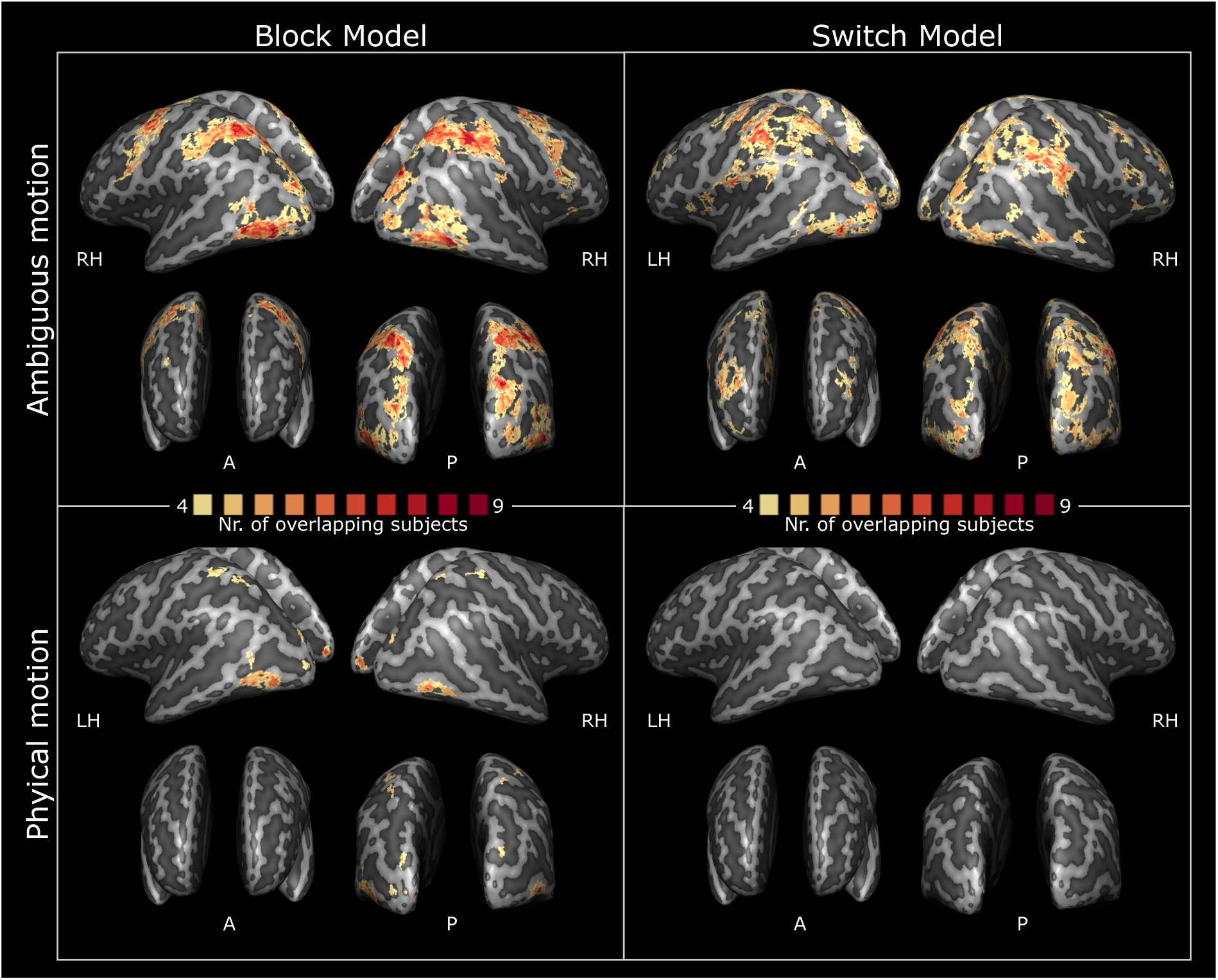
Conjunction of participants for the block and the switch model. Conjunction maps are generated by fitting the block model and the switch model to both the ambiguous and the physical condition. Voxels shown in warm colors survived q(FDR) *<* 0.05 in at least 4 of 9 participants. Cluster size threshold is applied to the surface maps (area *>* 20 *mm*^2^). Maps are shown on the inflated white matter surfaces reconstructed for both the left and right hemispheres in MNI space (A = anterior, P = posterior, LH = left hemisphere, RH = right hemisphere).

**Figure 3:**
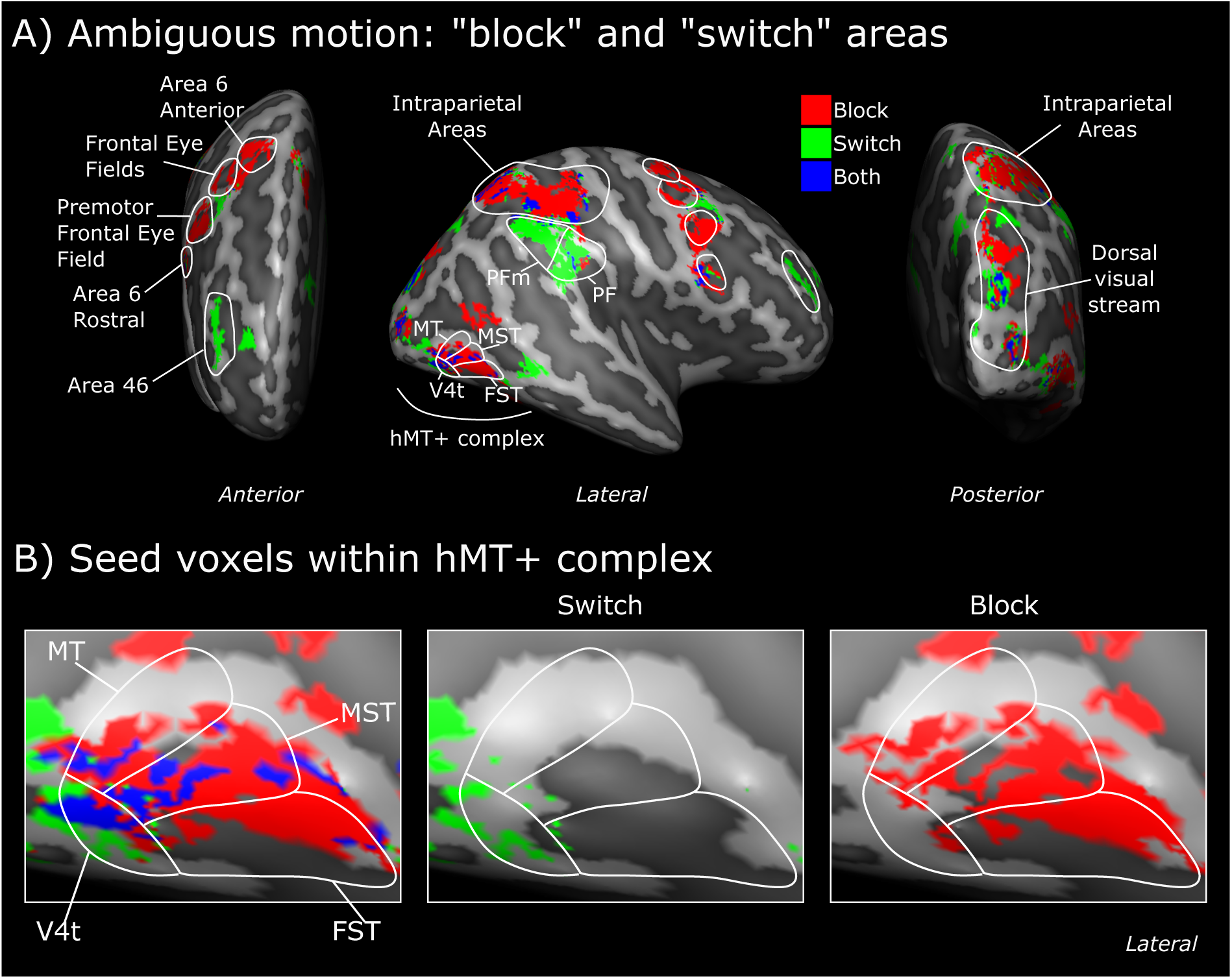
Group model-map for ambiguous motion condition. A) Conjunction maps are first thresholded by Nr *>* 4 and then converted into binary maps. The resulting binary maps, one for the switch and one for the block model are compared for each voxel and summarized in a categorical group map: a red label indicates the block model, a green label indicates the switch model and the blue color indicates that both models are passing the threshold. This categorical map is shown on the inflated white matter surface reconstructed for the right hemisphere in the MNI space. White lines encapsulating the resulting clusters of voxels indicate areal borders according to Glasser parcellation (Glasser et al., 2016). B) Voxels within hMT+ from the switch and the block models. The left panel is a zoom-in of the middle panel in A. The middle and right panels show the isolated switch and block voxels, respectively. Color coding is identical to A. Note that these switch and block voxels are used as separate seeds for subsequent connectivity analyses (Figure 4 and Figure 5).

**Figure 4:**
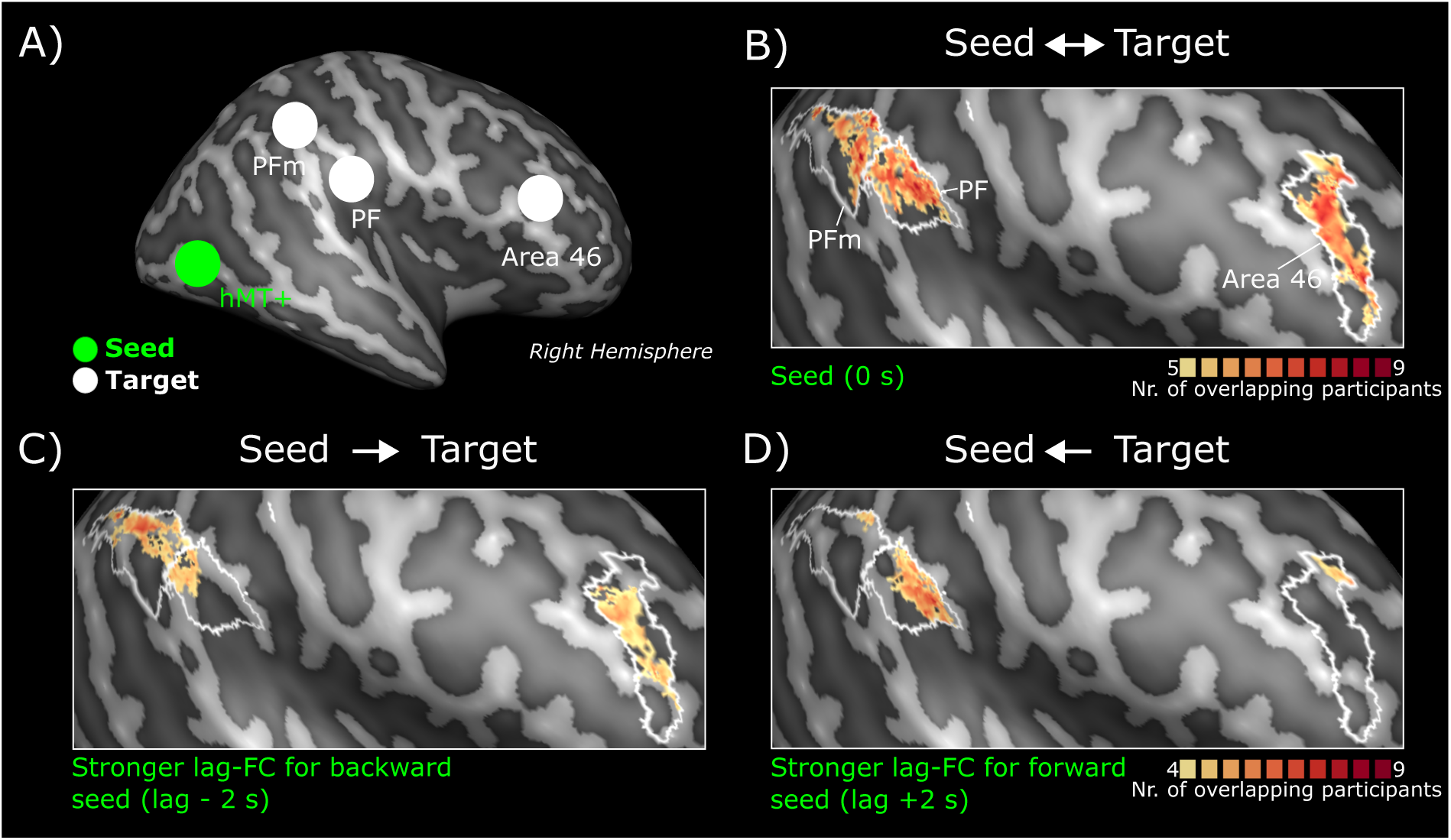
Lag-based and seed-based functional connectivity (FC) analysis for switch-type temporal dynamics . A) Schematic representation of the seed (green voxels within hMT+ indicated by the green circle) and the target areas (indicated by white circles). B) Group conjunction map resulting from voxel-wise correlation analysis (instantaneous synchrony, i.e. lag = 0). C) Seed-to-target group conjunction map. Lag-based FC for backward seed (lag = -2) *>* Lag-based FC for forward seed (lag = +2). D) Target-to-seed group conjunction map. Lag-based FC for backward seed (lag = -2) *<* Lag-based FC for forward seed (lag = +2)

**Figure 5:**
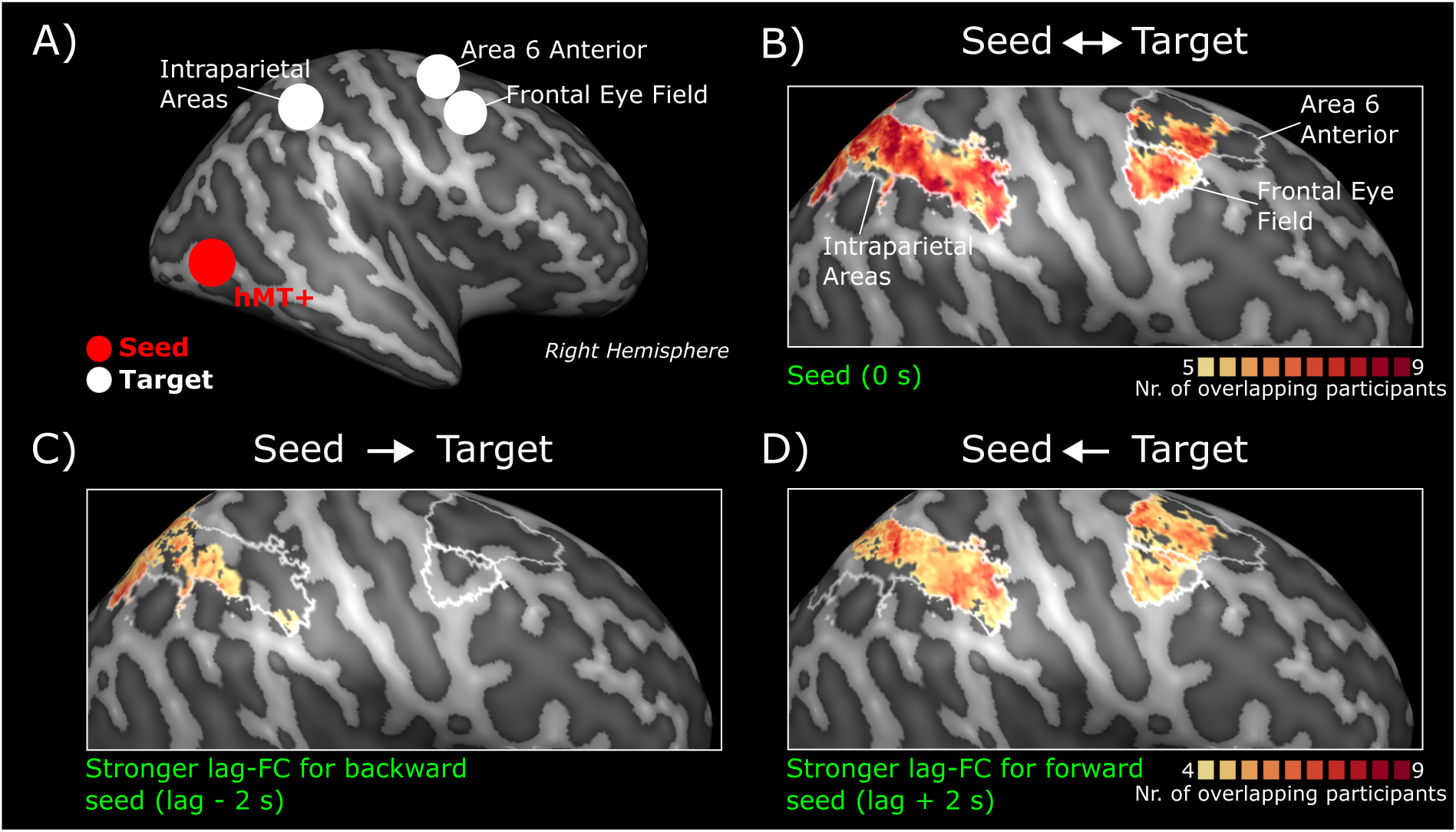
Lag-based and seed-based functional connectivity analysis for block-type temporal dynamics. A) Schematic representation of the seed (red voxels within hMT+ indicated by the red circle) and the target areas (indicated by white circles). B) Group conjunction map resulting from correlation analysis (instantaneous synchrony). C) Seed-to-target group conjunction map. Lag-based FC for backward seed (lag = -2) *>* Lag-based FC for forward seed (lag = +2). D) Target-to-seed group conjunction map. Lag-based FC for backward seed (lag = -2) *<* Lag-based FC for forward seed (lag = +2).

**Figure 6:**
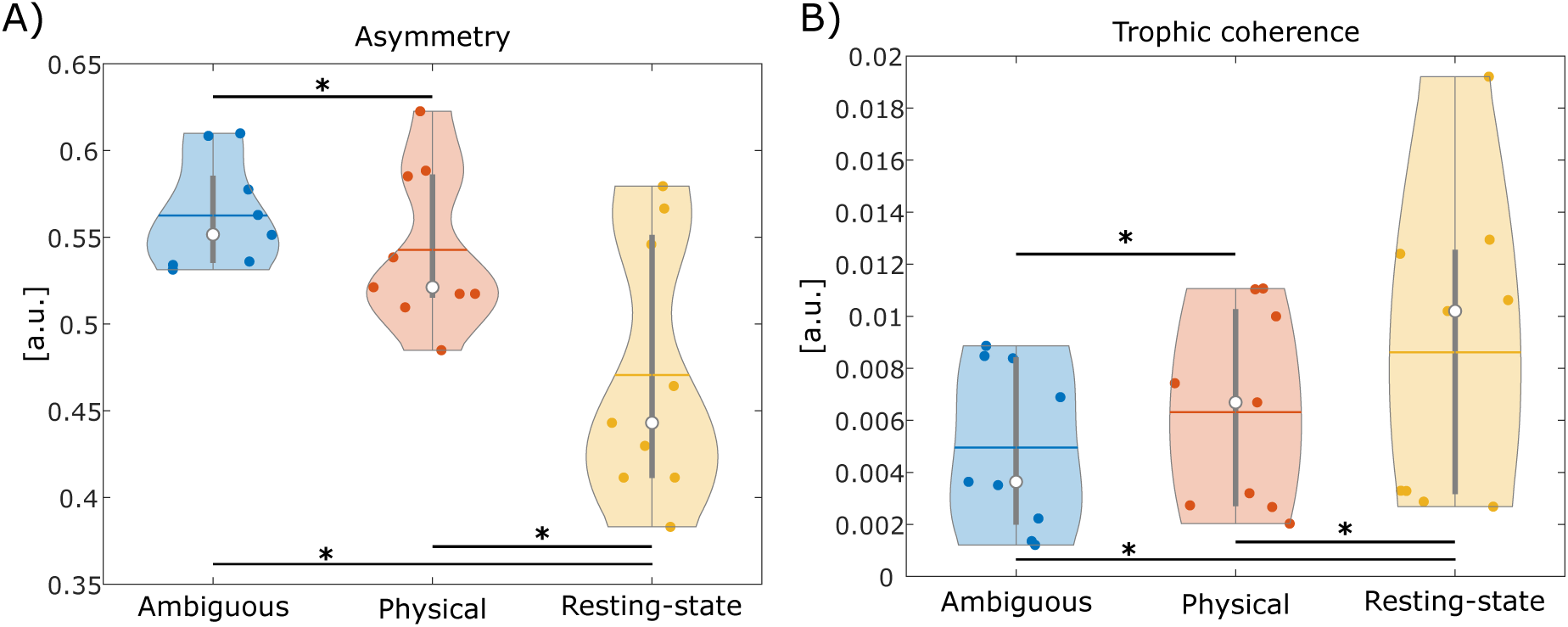
Distribution of asymmetry and trophic coherence in three brain states: ambiguous motion (task), physical motion (task) and resting state conditions. Each subject is represented as a dot within each distribution. Asterisks indicate statistical significance (t-test, FDR corrected) for p *<* 0.05.

**Figure 7:**
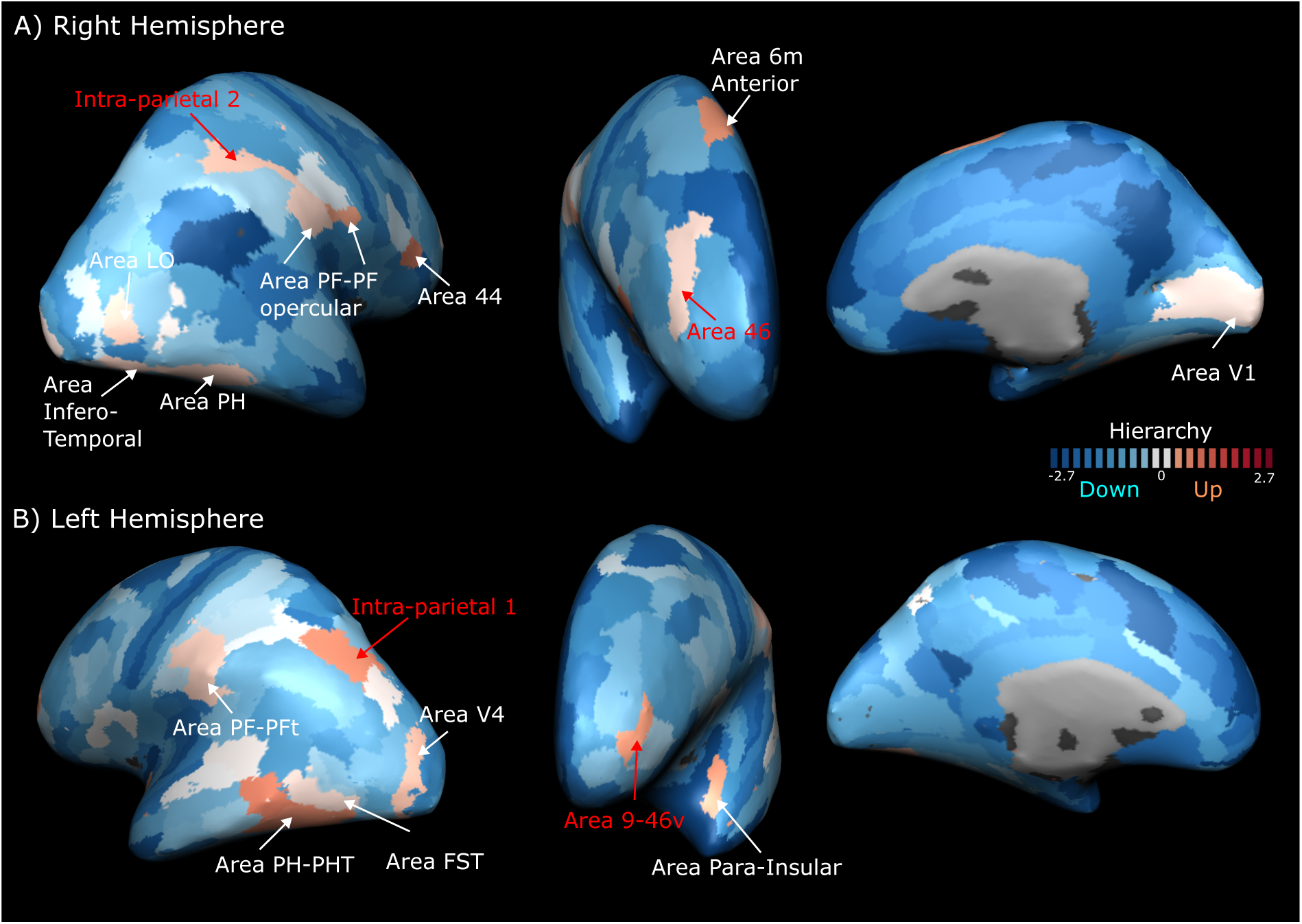
Changes in areas’ centrality from the physical to the ambiguous motion condition. Each area (Glasser parcellation, Glasser et al., 2016) is assigned a color representing the magnitude and direction of the centrality changes, averaged across participants. The red color bar indicates an increase in centrality, while the blue color bar indicates a decrease. White represents areas where the change is below 0.1 (excluded from further analyses).

## 2 Results

### 2.1 Brain responses to the ambiguous motion condition are captured by both the “block” and the “switch” models

During the ambiguous motion condition, participants alternate between perceiving two distinct perceptual states (horizontal and vertical motion) while the visual input remains constant. We hypothesized that these transitions between perceptual states and the periods of sustained perception would be reflected in neural activity with distinct temporal dynamics.To disentangle neural responses associated with sustained perceptual states from those linked to perceptual transitions, we implemented two general linear models (GLMs). The “block” GLM modeled the two perceptual states as sustained blocks to capture prolonged perceptual responses, while the “switch” GLM included transient predictors to model neural activity at moments of perceptual transitions. To investigate the differences between exogenously and endogenously driven perceptual transitions, we applied the same GLMs to both the physical motion condition, where perception is externally driven, and the ambiguous motion condition, where perception arises internally. This approach allowed us to directly compare the neural mechanisms underlying these two types of perceptual transitions. Our analysis revealed that, in the ambiguous motion condition, both block and switch predictors model the neural responses, consistent with the hypothesis that different temporal dynamics underlie sustained and transition-related mechanisms (**Figure 2**). However, the switch predictor did not explain neural activity during the physical motion condition, where responses were exclusively modeled by the block predictor. For the ambiguous condition, results from both models revealed consistent activation across participants, spanning much of the cortical landscape (**Figure 2**, first row). In contrast, for the physical condition, consistent activation across participants was only observed with the block model. Brain regions are labeled according to the Glasser parcellation (Glasser et al., 2016). Functional activations were predominantly within visual areas, including bilateral activation in V1, motion-sensitive regions (V4t, FST) in the human motion complex (hMT+), and sparse activation in dorsal and ventral visual areas extending to the parietal cortex. These findings indicate that neural modulation related to perceptual switches emerges specifically during the ambiguous motion condition, characterized by transient dynamics, whereas sustained perceptual responses are captured by prolonged activity during both the physical and the ambiguous condition. Unlike physical motion, where transitions are externally driven, ambiguous motion relies on endogenous processes during resolution of state transitions. As a result, neural populations encoding perceptual switches become functionally relevant only in the ambiguous condition. Moreover, the broader cortical engagement observed during ambiguous perceptual transitions aligns with previous reports linking higher-order cortical areas to bistable perception (J. Brascamp et al., 2018. This extended neural response underscores a distinctive aspect of human cognition, emphasizing the brain’s ability to resolve perceptual ambiguity through intrinsic processes.

### 2.2 Ambiguous motion dynamically modulates hMT+, frontal area 46, and parietal areas PM and PFm during the ambiguous condition

We summarize the group conjunction maps from our modeling analysis for the ambiguous condition (**Figure 2**, first row) by attributing to each voxel a label indicating the winning type of temporal dynamic: a block-like, a switch-like or mixed dynamics (**Figure 3**, **Table 1**). Within a Glasser cortical parcellation (Glasser et al., 2016), we only included areas that are consistently found in both hemispheres (**Figure 3**, circles and arrows point to bilaterally activated areas in the right hemisphere). This classification is particularly useful to understand how the brain organization changes during the ambiguous condition and how the temporal dynamic of each area might reflect the functional role in resolving the task. However, our results show that there are brain areas for which all voxels show the same temporal dynamics, but there are also areas in which subcategories of voxels can be found sharing different dynamics. For example, the human motion complex (MT, MST, V4t, FST) exhibit all three types of modulations. However, only MT, V4t and FST are bilaterally modulated in our group analysis. It is important to note that these distinctions are based on atlas-defined parcellation rather than functional localization and should therefore be considered approximate. Although small patches within MT and FST are found to be modulated by the switch model, the predominant ones show a block-like modulation. Differently, the voxels comprising area V4 transitional zone (V4t, Glasser area 156) are subdivided between the three types of modulation. This is different from the physical motion where FST and V4t show only a block-like modulation. The fact that hMT+ is involved in both stimulus types (physical and ambiguous motion) confirms previous fMRI results (Goebel et al., 1998; Muckli et al., 2002; Pizzuti et al., 2024; Schneider et al., 2019). However, an internal functional reorganization of subregions of the motion sensitive areas is expected to process and resolve a more complex task such as the ambiguous condition. Our results show that subdivisions of hMT+ might have different roles in resolving the ambiguity of the stimulus, MT and FST responsible for maintaining the perception during the sustained periods and that V4t responsible for the transitions between perceptual states and the sustained periods. This is in line with previous results investigating the role of subdivisions of hMT+ during complex tasks involving retinal and extraretinal signals (Sulpizio et al., 2022). Higher-order visual areas exhibit a similar mix of temporal dynamics as hMT+ and are grouped as ‘dorsal visual stream’ in **Figure 3**. Notably, during perceptual transitions, additional activation was observed in frontal areas 46, PFm, and the PF complex in the inferior parietal lobe (**Figure 3**, green color). Sustained perception, however, was associated with intraparietal areas (the Glasser parcels within this categorization are reported in **Table 1**, Panel A, ‘Parietal’ category in bold characters) along the superior parietal lobe, area 6 (anterior), the frontal eye field (FEF) which predominantly displayed block-like dynamics. These regions have previously been identified as neural correlates of bistable processing (J. Brascamp et al., 2018). Our analysis, however, distinguishes the functional contributions of areas within the fronto-parietal network, revealing a finer organization that supports the distinct phases of bistable perception: perceptual transitions and sustained perception.

**Table 1:** List of brain areas from Glasser atlas (Glasser et al., 2016) that are consistently found in at least 4/9 participants for both the block-model (A) and the switch-model (B). An area is included in this table if more than 5% of the total amount of voxels comprining the parcel is showing the same effect. Areas are divided in four macro-areas (frontal, parietal, temporal and occipital) according to their cortical location. For each entry, we report the number of the parcel (1-1180), the associated nome from Glasser parcellation (first letter indicates the hemisphere) and the number of voxels we found consistently modulated across subjects. In bold, areas bilaterally activated are highlighted.

| A) Block-model |  |  |  |
| --- | --- | --- | --- |
| FRONTAL | PARIETAL | TEMPORAL | OCCIPITAL |
| **10 L_Frontal_Eye_Fields, 43** | 17 L_IntraParietal_Sulcus_Area_1 | **23 L_Middle_Temporal_Area, 12** | **16 L_Seventh_Visual_Area, 17** |
| **96 L_Area_6_anterior, 87** | **48 L_Area_Lateral_IntraParietal_ventral, 77** | **156 L_Area_V4t, 86** | **21 L_Area_Lateral_Occipital_2, 40** |
| **1010 R_Frontal_Eye_Fields, 12** | **49 L_Ventral_IntraParietal_Complex, 36** | **157 L_Area_FST, 72** | **1016 R_Seventh_Visual_Area, 53** |
| 1011 R_Premotor_Eye_Fields, 59 | **50 L_Medial_IntraParietal_Area, 74** | 1002 R_Medial_Superior_Temporal_Area, 15 | 1019 R_Area_V3B, 23 |
| 1043 R_Supplementary_And_Cingulate_Eye, 43 | **95 L_Area_Lateral_IntraParietal_dorsal, 43** | **1023 R_Middle_Temporal_Area, 11** | 1020 R_Area_Lateral_Occipital_1, 11 |
| 1078 R_Rostral_Area_6, 44 | **117 L_Anterior_IntraParietal_Area, 48** | **1156 R_Area_V4t, 59** | **1021 R_Area_Lateral_Occipital_2, 28** |
| **1096 R_Area_6_anterior, 88** | **144 L_Area_IntraParietal_2, 72** | **1157 R_Area_FST, 68** | 1152 R_Area_V6A, 23 |
|  | 146 L_Area_IntraParietal_0, 23 |  | 1159 R_Area_Lateral_Occipital_3, 15 |
|  | 1046 R_Lateral_Area_7P, 18 |  |  |
|  | **1048 R_Area_Lateral_IntraParietal_ventral, 108** |  |  |
|  | **1049 R_Ventral_IntraParietal_Complex, 69** |  |  |
|  | **1050 R_Medial_IntraParietal_Area, 86** |  |  |
|  | **1095 R_Area_Lateral_IntraParietal_dorsal, 51** |  |  |
|  | **1117 R_Anterior_IntraParietal_Area, 135** |  |  |
|  | **1144 R_Area_IntraParietal_2, 40** |  |  |
| B) Switch-model |  |  |  |
| FRONTAL | PARIETAL | TEMPORAL | OCCIPITAL |
| 43 L_Supplementary_And_Cingulate_Eye, 39 | 14 L_RetroSplenial_Complex, 46 | 107 L_Area_TA2, 18 | 1 L_Primary_Visual_Cortex, 17 |
| 51 L_Area_1, 91 | 29 L_Medial_Area_7P, 24 | 175 L_Auditory_4_Complex, 50 |  |
| 54 L_Dorsal_area_6, 41 | 45 L_Medial_Area_7A, 58 | 1022 R_Posterior_InferoTemporal, 23 |  |
| 55 L_Area_6mp, 49 | 47 L_Area_7PC, 98 | 1025 R_Perisylvian_Language_Area, 54 |  |
| 57 L_Area_Posterior_24_prime, 25 | 117 L_Anterior_IntraParietal_Area, 52 | 1028 R_Superior_Temporal_Visual_Area, 27 |  |
| 113 L_Frontal_Opercular_Area_1, 12 | **144 L_Area_IntraParietal_2, 51** | 1047 R_Area_7PC, 50 |  |
| **84 L_Area_46, 17** | 147 L_Area_PF_opercular, 44 |  |  |
| 115 L_Frontal_Opercular_Area_2, 12 | 148 L_Area_PF_Complex, 69 |  |  |
| 59 R_Anterior_24_prime, 29 | **149 L_Area_PFm_Complex, 98** |  |  |
| **1084 R_Area_46, 77** | 1017 R_IntraParietal_Sulcus_Area_1, 14 |  |  |
| 86 R_Area_9-46d, 103 | 1095 R_Area_Lateral_IntraParietal_dorsal, 11 |  |  |
|  | **1144 R_Area_IntraParietal_2, 39** |  |  |
|  | **1149 R_Area_PFm_Complex, 238** |  |  |

### 2.3 Lag-based functional connectivity analysis reveal different functional coupling for perceptual transitions and sustained percepts during ambiguous motion

After identifying the main brain areas that are involved during the two phases (perceptual transitions and sustained perception) of the ambiguous motion condition, we turned our attention to investigating how these areas are functionally coupled in terms of effective connectivity estimates. For this purpose, we employed lag-based functional connectivity analyses (Deco et al., 2022) using the group hMT+ switch and block voxels as seeds (**Figure 3**B). For both seeds we estimated functional connectivity in each experimental condition across three distinct temporal lags (0,1, 2 s). Connectivity analyses on the ambiguous condition (with a lag of 2 s) were of main interest (**Figure 4**, **Figure 5**). Complementary analyses on the physical motion and resting state data, as well as the other lags, served as control (**Supplementary Figure 2 - 7**.

#### Perceptual transitions: frontal area 46 and PFm appear to predominantly follow the dynamics of hMT+

For the switch dynamics, we focus on the coupling between switch-based voxels within hMT+ and PF-PFm and frontal area 46 (**Figure 4A**). First, we found a significant instantaneous functional coupling between our seed and the targets (**Figure 4B**). This was expected as both seed and target are part of an extended network activated during perceptual switches. Second, our results on directionality estimates show that hMT+ seed seems to influence and drive the temporal dynamic of the frontal area 46 and the posterior area PFm. However, a reversed gradual trend towards the anterior part of the PF area is also observed (**Figure 4C-D**). In this simplified framework, our results might indicate that on one side hMT+ is following the dynamics of PF, on the other side driving Area 46 and part of PFm. In line with previous results, all three target areas have been reported as contributors of perceptual transitions (J. Brascamp et al., 2018). In addition, we now attribute to them a putative causal information or directionality of the information flow with respect to seed voxels within hMT+. Finally, the involvement of the dorsolateral prefrontal cortex (dlPFC), where area 46 is located remains a topic of debate during perceptual switches. Some studies have suggested its activation might be more related to increase of attention for the act of reporting perception rather than the switching process itself. Our results align with this hypothesis, as area 46 appears to follow the dynamics of hMT+, likely reflecting the temporal delay between the endogenous switch and the subsequent action of reporting the perception via button press.

#### Sustained perception: intraparietal areas and FEF appear to predominantly drive the dynamics of hMT+

For the sustained dynamics, we focus on the coupling between block-based voxels within hMT+ and intraparietal areas, frontal eye field and the anterior subregion of area 6 (**Figure 5A**). First, we found a significant instantaneous functional coupling between our seed FEF and part of area 6 anterior (**Figure 5B**). Second, our results on directionality estimates show that our hMT+ seed seems to differently interact with the targets: while the hMT+ seed seems to influence and drive the temporal dynamic of the posterior part of the intraparietal areas, the more anterior part of the intraparietal areas, FEF, and area 6 anterior seem to drive the dynamic of the hMT+ seed (**Figure 5C-D**). Similarly as for the perceptual transition case, all target areas have been reported as contributors of solving bistable stimuli (J. Brascamp et al., 2018). A high level of specialization of these target areas is also expected from bistable human perception literature. For example, Knapen and colleagues found that differential specialization of intraparietal areas compared to FEF and surrounding areas according to the type of duration of the perceptual transitions during binocular rivalry tasks (Knapen et al., 2011).

### 2.4 Enhanced asymmetry and reduced feedback loops characterize the perception during the ambiguous motion condition

We hypothesized that global quantitative measures of brain hierarchical organization, such as asymmetry and trophic coherence, could reveal distinct functional organizations underlying three brain states: the ambiguous motion condition, the physical motion condition, and the resting-state condition. These measures were derived from the generative effective connectivity (GEC) matrix obtained by fitting empirical fMRI time series to the Generative Connectivity of the Arrow of Time (GCAT) model. At the group level, we observed significant trends in both asymmetry and trophic coherence across the three states (q(FDR) *<* 0.05; **Figure 6**). Specifically, asymmetry decreased from task conditions to resting state, while trophic coherence exhibited the opposite trend, increasing from tasks to rest. Notably, the ambiguous motion condition showed the highest asymmetry and the lowest trophic coherence, distinguishing it from the other states. These results suggest that the complexity of the task is reflected in these global network measures. Asymmetry quantifies the directional flow of information between pairs of brain regions across the network (M. L. Kringelbach et al., 2024). As previously reported, higher asymmetry during tasks reflects the more complex computations required for task conditions compared to resting state (Deco et al., 2022). In contrast, trophic coherence measures the directness of the network in terms of feedback loops, providing an indication of network stability: lower trophic coherence reflects fewer feedback loops and corresponds to higher network stability (Dambrot, 2017a). In conclusion, the resolution of an ambiguous stimulus manifests itself in a network with greater asymmetry, leading to a more hierarchical organization. This hierarchy, in turn, enhances network stability, as indicated by lower trophic coherence, with fewer feedback loops needed to maintain functionality.

### 2.5 Changes in areas’ centrality from the physical to the ambiguous motion condition

As our results indicate a distinct orchestration of brain activity during the ambiguous motion condition compared to the physical motion condition (**Figure 6**), we further investigated whether specific areas drive this change. For both motion conditions, we computed the total degree (*G tot*) for each node, representing its centrality within the network. As previously proposed, this centrality measure is also related to a measure of generative hierarchy within the network (M. L. Kringelbach et al., 2024. Our analysis revealed that the centrality of the majority of brain areas changes to accommodate the differing task demands (**Figure 7**). Specifically, from the physical to the ambiguous condition, the centrality of most nodes decreases, a shift counterbalanced by a subset of areas showing increased centrality. These areas include: right frontal area 46 and left area 9-49, bilateral area PF and its adjacent right OF opercular, left intraparietal areas 1 and 2, right area 44 and right anterior area 6, right inferotemporal area, right area LO and V1, bilateral area PH, left area PHT, left area FST, left area V4, and the left para-insular area. These regions appear critical for the brain’s network organization during the ambiguous condition. To interpret these findings in functional terms, we incorporated insights from the temporal dynamics characterized by the switch and block models at the voxel-wise level (**Figure 2**). For instance, frontal area 46 and area PF, which were associated with the switch model, suggest a pivotal role for the network during perceptual transitions. Conversely, intraparietal areas 1 and 2, linked to the block model, imply a key involvement in the network during sustained perception. These results highlight the dynamic reorganization of brain network centrality as a mechanism for resolving ambiguous stimuli, emphasizing distinct roles for specific regions during perceptual transitions and sustained perception.

## 3 Discussion

### 3.1 Summary

In this whole-brain fMRI study, we used the ambiguous motion quartet stimulus, which elicits bistable motion perception and induces a series of endogenous perceptual transitions between two motion states (vertical and horizontal) under constant sensory input. While the involvement of parietal and frontal areas in resolving ambiguity has been widely discussed in the literature (J. Brascamp et al., 2018), their specific functional roles remain under debate. To address this, we combined voxel-wise analysis (GLM), lag-based functional connectivity (lag-based FC) analysis, and a whole-brain computational model of generative effective connectivity (GCAT). The GLM approach disentangles brain activity associated with perceptual transitions from that related to sustained perception, while the lag-based FC and GCAT analyses investigate directionality of the information flow between brain areas and changes in hierarchical brain organization. Together, these approaches helped identify key regions driving hierarchical processes during ambiguous versus physical motion conditions (control condition).

### 3.2 Perceptual transitions and sustained perceptual states

Our study provides a nuanced perspective on the neural mechanisms underlying bistable motion perception by explicitly distinguishing between perceptual transitions and sustained perceptual states during the ambiguous motion condition. Regions activated by the switch model (**Figure 3**, green voxels) closely align with findings from prior studies, as comprehensively reviewed by J. Brascamp et al. (2018). However, in our case, key areas, such as the frontal eye fields (FEF), intraparietal regions within the superior parietal lobule (SPL), and hMT+, were better captured by the block model compared to the switch model. This phase of bistable processing, the sustained state, has received less attention compared to transitions, and only a few studies have specifically addressed it. Notably, Muckli et al. (2002), using apparent motion stimuli with fMRI, revealed that hMT+ contributes more to the sustained phase than to the transition. In our study, the majority of hMT+ voxels were captured by the block model, although a small subset also showed modulation consistent with the switch model. This finer organization of hMT+ might stem from the increased sensitivity and spatial resolution afforded by 7T imaging in our study. Knapen and colleagues investigated neural bistability using apparent motion and binocular rivalry with fMRI in humans looking at both switch and sustained responses, but they reported no differences between these two (Knapen et al., 2011). However, their stimuli did not evoke sharp, instantaneous transitions like our ambiguous motion quartet. Instead, they emphasized the role of transition duration, offering a more refined characterization of fronto-parietal network areas. These findings suggest that regions within this network may exhibit functional flexibility, with subregions adapting to the specific demands of bistable perception tasks.

### 3.3 The role of the right frontal area 46 in perceptual transitions

Consistent with previous findings on bistable paradigms (J. Brascamp et al., 2018), we observed that the frontal area 46 is associated with perceptual transitions and predominantly lateralized in the right hemisphere. This right-lateralization aligns with the conventional explanation that bistable processing is closely tied to attention (Lumer et al., 1998), where the dominance of parietal areas for spatial attention (Corbetta and Shulman, 2002; Sheremata and Silver, 2015) and the distribution of noradrenergic locus coeruleus terminals to frontoparietal regions (Corbetta et al., 2008) play central roles. These right-lateralized regions are thought to mediate increases in arousal, alertness, and focused attention, facilitating perceptual transitions. However, more recent theories suggest that this right-lateralization of frontal areas is a consequence of the increased attention that is not needed for the transition itself but for its report with a motor action (J. Brascamp et al., 2018). In our experimental paradigm, participants reported their perceptual transitions during the fMRI session by pressing a button with their right hand. While we can exclude a strict somatosensory circuit association (as this action correlates with somatosensory/motor areas in the left-hemisphere), we cannot rule out whether the source of the activation of frontal area 46 is related to the action of the report. Indeed, studies using non-report paradigms, such as J. W. Brascamp et al. (2015), showed reduced frontoparietal activation during bistable tasks, supporting the notion that this activation may be a consequence, rather than a cause, of perceptual switches (J. W. Brascamp et al., 2015). Our lag-based connectivity analysis supports this interpretation, revealing a directed flow of information from hMT+ to frontal area 46. If hMT+ resolves stimulus ambiguity and initiates motion perception (Salzman et al., 1992; Schneider et al., 2019), a temporal delay is expected between the switch and the subsequent reports through motion action. According to this theory, the activation of frontal area 46 is expected to follow hMT+ and to reflect the increased attention prior to the conscious report. To clarify this further, future studies could employ non-report paradigms to decode perceptual switches during ambiguous motion tasks. For example, eye-tracking (Frässle et al., 2014; Naber et al., 2011) or decoding perceptual states directly from fMRI data could provide further insights. Additionally, transcranial magnetic stimulation could be applied to the right frontal area 46 to examine its causal role in perceptual switches. While such TMS methods have been widely used to explore the function of parietal areas, their application to frontal areas remains comparatively underexplored.

### 3.4 Directionality and hierarchy estimates for complex cognitive tasks

The application of lag-based functional connectivity (lag-based FC) to estimate the direction of information flow between brain areas using fMRI time series (Figures 4-5) draws inspiration from the ‘inside-out’ framework (Deco et al., 2022). In this framework, lag-based FC has been used as a data-driven approach to successfully infer the asymmetry of information flow and the arrow of time for different cognitive tasks (Deco, Perl, et al., 2021; Deco et al., 2023; Guzmán et al., 2023; M. Kringelbach et al., 2022; M. L. Kringelbach et al., 2024). Among the realm of methods to estimate effective connectivity (Friston, 2011; M. L. Kringelbach et al., 2024), lag-based FC can be considered a simplified alternative to Granger Causality (GC), which has been successfully applied to fMRI for estimating directionality (Goebel et al., 2003; Roebroeck et al., 2005). Unlike GC, which requires constructing autoregressive models for both the seed and the target regions, lag-based FC significantly reduces computational demands by calculating only two correlation coefficients (comparing the target signal with two shifted versions of the seed time course). Moreover, lag-based FC has proven advantageous for enhancing the performance of generative whole-brain models of effective connectivity (Deco et al., 2024; Gilson et al., 2016; M. L. Kringelbach et al., 2024, a model also utilized in this study. Given these benefits and the expected similarity in outcomes between lag-based FC and GC, we opted for the former in our analysis. Future investigations might complement our findings with results from GC and investigate whether more detail in the computations is required to extend our understanding of the phenomena. Alternatively, dynamic causal modeling (Friston et al., 2003, 2019; Stephan et al., 2010) offers a robust approach for further validating our results on lag-based functional connectivity. Specifically, given that the number of regions involved during both the switch and sustained phases is relatively small, we could perform Bayesian model comparison of competing hypotheses to test the two directions of information flow to and from hMT+ in each phase. Finally, as a further extension of data-driven measure of directionality between couples of brain areas, we used the GCAT modeling approach and explored the link between directionality and changes in hierarchical organization. Consistent with prior findings, network asymmetry and trophic coherence measures (Deco et al., 2024; M. Kringelbach et al., 2022) captured the complexity of brain states (**Figure 6**). In addition, when comparing the brain hierarchical organization between the ambiguous and the physical condition (**Figure 7**) in terms of node’s centrality, a proportion of the resulting brain areas (frontal area 46, intraparietal area 1, parietal area PFm) from this analysis coincide with the areas we detected using simple general linear modeling (**Figure 3**). While this series of results further highlight the involvement of these brain areas during the ambiguous motion task, it’s still unknown whether the presence of these brain areas is “essential” for the network for resolving the cognitive task. Recent computational models have begun to investigate the effects of perturbing or disconnecting brain areas to study their impact on network dynamics (Cabral et al., 2012; Deco et al., 2018; Idesis et al., 2024; Thiebaut de Schotten et al., 2020). While originally applied to model brain configurations in pathological conditions, these approaches hold great potential for advancing our understanding of human cognition in healthy states.

### 3.5 Conclusions

To conclude, our findings provide new insights into the neural mechanisms underlying bistable perception, highlighting the dynamic interplay between brain regions during transitions and sustained perception. Specifically, we show that during the perceptual switch, hMT+ interacts with frontal (area 46) and parietal areas PM and PFm, while during sustained perception, it engages with intraparietal regions. By integrating voxel-wise analysis, lag-based functional connectivity, and generative computational modeling, we demonstrate that perceptual ambiguity elicits reorganization of hierarchical networks, with frontal and parietal areas ascending in functional hierarchy to coordinate ambiguity resolution. These results underscore the adaptability of the brain’s hierarchical organization in meeting the demands of complex cognitive tasks, paving the way for future studies to explore causal manipulations of these networks and their role in shaping perception.

## 4 Materials and methods

### 4.1 Experimental design

#### 4.1.1 Ethics

Written informed consent was obtained from each participant before the experiment began. The study was approved by the ethics review committee of the Faculty of Psychology and Neuroscience (ERCPN) of Maastricht University and experimental procedures followed the principles expressed in the Declaration of Helsinki.

#### 4.1.2 Participants

Nine healthy individuals (7 females and 2 males) with normal or corrected-to-normal vision participated in the study. All participants were compensated financially for their involvement and had prior experience with MRI scanning. The study comprises one hour behavior session (participant selection according to behavioral measure) and a 2 hours fMRI session at 7 T. The behavioral pre-scan session aimed to account for the expected variability in individual responses to the ambiguous motion stimulus (Pizzuti et al., 2024; Schneider et al., 2019). Participants who exhibited a stable alternation between horizontal and vertical motion (or vice versa) at intervals of at least 5 seconds on average, evaluated over three runs of ambiguous motion stimulus in a behavioral laboratory setting, were selected for the fMRI session.

#### 4.1.3 Stimulus description

We used the same motion quartet stimulus used in our previous study (Pizzuti et al., 2024) to evoke both real and illusory horizontal and vertical motion. The motion quartet is characterized by four square “inducers” (1° × 1° visual angle) arranged with a 3° horizontal and 3.7° vertical distance (Pizzuti et al., 2024; Schneider et al., 2019). In the physical quartet, squares physically moved horizontally and vertically for 10 s per motion type. After four motion alternations (80 s total), a 16 s flicker condition, where all inducers blinked synchronously, served as a baseline without inducing motion. This sequence is repeated six times, with runs beginning and ending with 19-21 s of fixation. For the ambiguous quartet, the horizontal and vertical motions were replaced with a constant ambiguous stimulus (80 s) that induced apparent horizontal or vertical motion through alternately blinking diagonal squares. Each square pair was displayed for 150 ms (9 frames) with a 67 ms inter-stimulus interval (4 frames), a presentation frequency (2.3 Hz) known to induce apparent motion (Finlay and von Grünau, 1987; Pizzuti et al., 2024; Schneider et al., 2019). While in the scanner, participants reported their percepts (horizontal or vertical) using an MR-compatible button box. Each physical and ambiguous run lasted 10 min 20 s (616 volumes, TR = 1 s). Finally, we collected one run of resting-state (rs-fMRI) lasting 10 min (600 volumes, TR=1 s), while the participant was asked to fixate a black fixation cross on a gray background. A frosted screen (distance from eye to screen: 99 cm; image width: 28 cm; image height: 17.5 cm) at the rear of the magnet was used to project the visual stimuli (using Panasonic projector 28 PT-EZ570; Newark, NJ, USA; resolution 1920×1200; nominal refresh rate: 60 Hz) that participants can watch through a tilted mirror attached to the head coil. We used 50% gray background (at 435 cd/m2 luminance) with white dots (at 1310 cd/m2) for the motion stimulation (black color is measured at 2.20 cd/m2). The scripts used for the stimulus presentation were developed in PsychoPy3 (v2020.2.4) and made available on Github https://github.com/27-apizzuti/macro_MotionQuartet.

### 4.2 Data acquisition and processing

fMRI data were collected with the whole-body MAGNETOM 7 T “Plus” (Siemens Healthineers, Erlangen, Germany) at Scannexus B.V. (Maastricht, The Netherlands) using a 32-channel RX head-coil (Nova Medical, Wilmington, MA, USA). fMRI data with whole-brain coverage were acquired using 2D GE EPI sequence with BOLD contrast (based on Moller2010) with the following imaging parameters: voxel resolution = 1.8×1.8×1.8 mm, echo repetition time (TR) = 1000 ms, echo time (TE) = 22 ms, nominal flip angle (FA) = 60°, GRAPPA = 2, multi band factor (MB) = 4, 72 slices, phase partial fourier =, field of view = 198 x 198 mm, data matrix: 110×110. Before the acquisition of the run, we collected 5 volumes for (offline) distortion correction with the settings specified above but opposite phase encoding (posterior-anterior). MP2RAGE (magnetization prepared 2 rapid gradient echoes) (Marques et al. 2010) at 0.7 mm isotropic resolution images were already available for all the participants from our previous experiment (Pizzuti et al., 2024).

fMRI data were preprocessed as follows: slice time correction (BrainVoyager v.22.4, later referred as ‘BV’, Goebel, 2012), motion correction (BV), geometric distortion correction (fsl-topup v.6.05, Smith et al., 2004), linear trend removal and high-pass filter (3 cycles for resting-state, 5 cycles for physical and ambiguous conditions using BV). The pair of opposite phase encoding images acquired at the beginning of the first functional run were used to estimate the susceptibility-induced off-resonance field and correct for geometric distortions. Boundary-based registration as implemented in BV was used to align fMRI data to T1-weighted anatomical data (T1w) from MP2RAGE. Anatomical T1w images were preprocessed as follows: upsample to 0.6 mm iso. (BV), skull stripped (fsl-bet), corrected for field inhomogeneity (N4biasFieldcorrection, Tustison et al., 2010 and registered to MNI space using linear and non linear co-registration algorithm ‘Syn’ from ANTs v.20.3.01 (Avants et al., 2008). We applied the set of transformation matrices for the MNI space to the functional data by using ANTS apply transformation command with Lanczos interpolation method. The spatial resolution of fMRI data was kept at original 1.8 mm iso. resolution. Once in the MNI space, we applied Glasser atlas to parcel the whole brain (Glasser et al., 2016). The Glasser parcellation scheme divides the brain into 360 nodes (180 nodes per hemisphere) and it is based on structural, functional, and connectivity features.

### 4.3 General linear modeling: the block and the switch model

While perceiving the ambiguous motion stimulus, the brain spontaneously and endogenously alternates between horizontal and vertical motion perception, even though the sensory input on the screen remains constant. This dynamic likely engages brain regions responsible for driving perceptual switches and others for sustaining the perception of motion. To distinguish these roles at the voxel level, we employed two general linear models (GLMs): the block-model and the switch-model. The block-model identifies sustained activity associated with perceiving horizontal or vertical motion, while the switch-model captures transient activity during transitions between these percepts. In the block-model, we modeled two predictors (horizontal motion and vertical motion) using canonical hemodynamic response functions (HRFs) with a two-gamma model. In the physical runs, these predictors were time-locked to the presentation of the stimuli, with each motion block lasting 10 seconds. For the ambiguous runs, predictors were time-locked to the subject’s reported perceptual state and varied in duration according to the subjective length of each perceptual block. In the switch-model, like the block-model, we included two predictors (horizontal motion and vertical motion) modeled with canonical HRFs. In the physical runs, these predictors were time-locked to the stimulus transitions, while in the ambiguous runs, they were time-locked to the subject’s reported perceptual switches. Unlike the block-model, the response duration in the switch-model was fixed at 1 second to specifically capture activity linked to perceptual transitions rather than sustained perception. Note that the physical condition serves as a control for the ambiguous condition, as motion switches are driven by the explicitly defined and stable physical motion stimuli. For both models, voxel time courses were normalized in BrainVoyager using the formula *y_n_orm* = *y/y_m_ean* ∗ 100, and GLMs were corrected for temporal autocorrelation (AR2). Each GLM runs for each subject and condition (physical and ambiguous) separately by concatenating runs of the same types. For each voxel, the estimated beta is then converted into a statistical t-value that tests whether the effect of both predictors (horizontal and vertical) is significant with respect to the baseline condition (flicker condition, not modeled as predictor in the GLM). Finally, the significance of the resulting t-maps for each subject and condition is corrected for multiple comparisons by using the false discovery rate (FDR) approach (Benjamini and Hochberg, 1995) considering the total number of voxels within a brainmask. As a result, each subject is characterized by four t-maps FDR corrected p *<* 0.05 (2 conditions x 2 GLM models).

### 4.4 Group conjunction maps

As all the functional maps are evaluated in the common MNI space, we investigated the spatial consistency of functional activity patterns across subjects by calculating probabilistic functional maps for both the block and the switch model evaluated for the physical and the ambiguous condition. Conjunction maps or probability maps (scaled in range 0-10) are a general method commonly used for quantifying spatial consistency of anatomical and functional areas to find locations of high consensus across a group (**Sitek2019**; Dworetsky et al., 2021; Gulban et al., 2020; Rosenke et al., 2021). At each spatial location, such maps represent the number of subjects with significant task activity (q(FDR) *<* 0.05) (results shown in **Figure 2**). Finally, to compare results from both the block and the switch model, we thresholded the respective conjunction map (group consistency above 55%, for a number of subjects *>* 4) and created a binary map (results from the block model were labeled with ‘1’, while results from the switch model were labeled with ‘2’) (results shown in **Figure 3**). The categorical map resulting by summing the block and the switch binary maps indicates which model (block, switch or both models) consistently captures the dynamic of each voxel.

## 5 Seed-based functional connectivity analysis

To investigate whether the motion complex (hMT+) is functionally connected with other brain areas during the ambiguous motion condition, we performed a seed-based functional connectivity analysis. We define two seeds of interest within the motion complex (Glasser areas MST, MT, V4t, FST; similar definition of hMT+ Huang et al., 2019; Kolster et al., 2010; Sulpizio et al., 2022): one hMT+ seed is composed by the voxels whose temporal dynamics were captured by the switch model and the other seed is composed by the voxels whose temporal dynamics were captured by the block model for the ambiguous motion condition. For each run and subject, the seed time course is created by averaging the selected voxel time courses over the two hemispheres. Then, a seed-based functional connectivity analysis is computed by calculating the Pearson correlation value (and its associated p-value) between the seed time course and every other voxel within the brain. This computation results in a correlation and a p-value map. Correlation is a well-known measure of synchrony or functional connectivity between two brain regions. For each subject and condition, we averaged the correlation maps and p-value maps across runs. Finally, we corrected the computed p-value maps for multiple comparisons using the FDR correction method (Benjamini and Hochberg, 1995). For this step, only voxels within a cortical brainmask are considered. Finally, we evaluated the spatial consistency of thresholded correlation maps (q(FDR) *<* 0.05) across participants by creating a group conjunction map (results shown in **Figure 4**, **Figure 5**).

## 6 Lag-based functional connectivity analysis

To explore the causal relationship between the motion complex (hMT+) and other brain areas, we employed a seed-based lag-based connectivity analysis. Shifted or lag-based correlations have been widely used to infer causal interactions between brain regions in fMRI studies, both in data-driven approaches (Deco et al., 2022) and in modeling frameworks (Gilson et al., 2016). Building on this foundation, we extended our connectivity analysis to incorporate this method. Specifically, the time course of a seed region of interest is shifted backward and forward by a specified time lag, and the Pearson correlation coefficient is calculated with respect to all voxels within a brain mask. The time lag is expressed as a multiple of the fMRI sampling rate (TR = 1 s). For each pair (seed and voxel time series), identifying the direction of the shift that produces the strongest correlation provides insights into causal dependencies between the variables. For example, for a time lag of 2 seconds, we computed two lag-based correlation maps: one for a positive lag of +2 s (shifting the seed time course forward) and another for a negative lag of -2 s (shifting it backward). The positive lag map reflects how strongly the seed “follows” other areas (occurring earlier in time), while the negative lag map indicates how strongly the seed “drives” activity in other brain areas. This method was applied to each subject for each experimental run across three conditions: ambiguous motion, physical motion, and resting-state. While the ambiguous condition was our primary focus, the physical and resting-state conditions served as controls. Additionally, lag-based correlation maps were computed for a range of lags (1, 2, and 3 seconds) by using the two hMT+ seeds as defined in the paragraph “Seed-based functional connectivity analysis” (**Figure 3**B). In order to choose the optimal range of lags for capturing causal information, we conducted an autocorrelation analysis on the fMRI time series as previously done (Deco et al., 2022; Guzmán et al., 2023). Briefly, we computed the autocorrelation for all regions, subjects and conditions, and chose the lag for which we observed a sufficient decay of the autocorrelation function (between 30-50%). Consistent with previous findings (Deco et al., 2022; Guzmán et al., 2023), we found that a sufficient decay occurs with lags = 1-3 s (**Figure 1**). Briefly, after computing the lag-based correlation and associated p-value maps for each run, we averaged these maps across runs within the same condition. The resulting p-value maps were corrected for multiple comparisons using the FDR correction method, considering only voxels within a brain mask. Finally, a single-subject difference map was generated by subtracting the correlation map for lag = -2 (seed driving) from the correlation map for lag = +2 (seed following). These difference maps were then thresholded and converted into binary maps, retaining the sign of the original difference: positive values were assigned +1, and negative values were assigned -1. At the group level, we created conjunction maps to identify voxels that consistently showed the same modulation with respect to the seed across subjects.

## 7 Whole brain generative effective connectivity modeling

While our empirical measure of connectivity focuses on local dynamics between the seed area (hMT+) and the rest of the brain, whole brain computational models offer a complementary perspective on the underlying global mechanisms and the hierarchical organization characterizing a brain state. Here, we construct a generative whole brain model according to the Generative Connectivity of the Arrow of time (GCAT) framework (M. Kringelbach et al., 2022) in which the brain is represented as a network of coupled linear Hopf oscillators. This method has been extensively used and successfully proved to replicate features of brain dynamics in both fMRI data (Deco, Vidaurre, and Kringelbach, 2021; M. Kringelbach et al., 2022; M. L. Kringelbach et al., 2015) as well as electrophysiology and magnetoencephalography data (Deco et al., 2017; Freyer et al., 2012). In our case, our brain network is constructed by 360 Hopf linear oscillators (Glasser parcellation) interconnected by a given coupling matrix. See Ponce-Alvarez and Deco (2024) for details on the linear approximation of the non-linear Hopf oscillator. The intrinsic frequency of each oscillator is computed from the data, as the averaged peak frequency of the BOLD signal of each brain region (M. Kringelbach et al., 2022). As the nodes constitute a network that generates whole-brain dynamics, the local dynamic of each node is described by a ‘revised’ version of the Landau-Stuart oscillator, that while modeling the transition from noisy to oscillatory dynamics, has in addition a coupling factor indicating that nodes are not isolated but influencing each others (Ponce-Alvarez and Deco, 2024, Equ. 9). The coupling information of the network, later referred to as generative effective connectivity (GEC), is initially attributed to structural connectivity (GEC=SC). We used the SC matrix for the Glasser parcellation that was made available by the authors of (Deco, Vidaurre, and Kringelbach, 2021). This SC was estimated from the standard HCP multi-shell dMRI data from 985 HCP participants (http://www.humanconnectome.org/), refer to Deco, Vidaurre, and Kringelbach (2021) for more details on processing steps. Through an interactive approach, this GEC matrix will be updated to fit the empirical features from the data until the optimization converges to a stable value. More specifically, the empirical features used to optimize the GEC are the functional connectivity matrix - FC (Pearson correlation) and time-lagged shifted covariance matrix - COVtau for a specified time shift (tau), that in our data was chosen tau = 2 s according to our autocorrelation analysis as explained in ‘Lag-based functional connectivity analysis’ paragraph.

During model optimization (Eq.1), the FC matrix and COVtau were updated according to epsFC = 0.0004 and epsCOVtau = 0.0001.

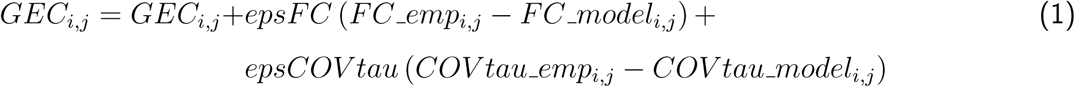

The resulting GEC is an asymmetric matrix with dimension (NxN, N = 360 nodes), where each entry quantifies the directed information flow between two brain regions. For example, *GEC_i,j_* quantifies the interaction between node i and node j, in terms of incoming level of information from node j to i. Contrary, *GEC_j,i_* quantifies the incoming level of information from node i to j. Leveraging the information contained in such a matrix, quantitative metrics can be quantified to describe the brain dynamics. A measure of node centrality is computed as total degree *G tot_i_* by quantifying for each node:

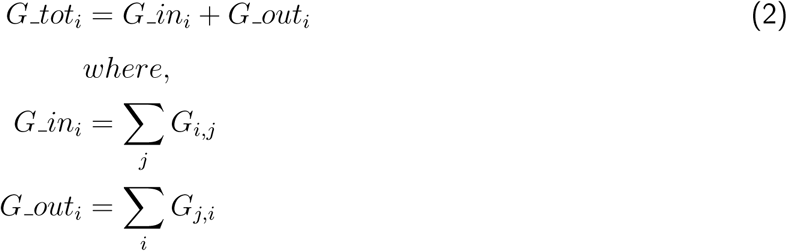

## 8 Asymmetry and trophic coherence measures

Once the GEC is estimated for a given brain state, due to its asymmetric nature, it can be used to infer characteristics of the hierarchical organization of the brain. It has been shown that when the brain is engaged in complex tasks, a stronger hierarchical organization appears compared to the resting-state condition as more computations are required in a task condition (M. Kringelbach et al., 2022). Stronger is the hierarchical organization, more asymmetric the GEC will be. In order to characterize the hierarchical organization of the three brain states elicited by the ambiguous, the physical and rest condition, we computed a measure of asymmetry of the GEC as number of asymmetric cells greater than a threshold (0.1) within the GEC (Eq. 3):

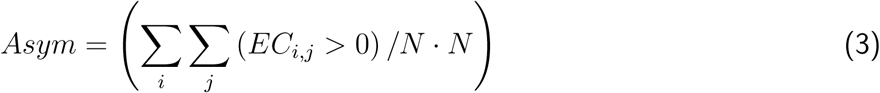

where *EC* = |*GEC* − *GEC^T^* | and N = 360 (number of nodes)

Complementary to a measure of the asymmetry, we also computed a measure of trophic coherence (TC) from the GEC matrix as previously done (Deco et al., 2024; M. L. Kringelbach et al., 2024). This measure from graph theory has been found to be relevant in describing a hierarchical network in terms of stability and feedback loop (Dambrot, 2017b; Mackay et al., 2020). In particular, the trophic coherence is an indication of the amount of feedback loop presented in the system: it has been demonstrated that a configuration of a hierarchical network characterized by a high stability is explained by the presence of few feedback loops. This is reflected in a low measure of TC.

## 9 Data and Software availability statement

Analysis code is available on GitHub: https://github.com/27-apizzuti/macro MotionQuartet. Raw fMRI data are available on Zenodo: https://doi.org/10.5281/zenodo.14809131.

## 10 Acknowledgements

This project was funded by the EU-project H2020-860563 euSNN and the European Union’s Horizon 2020 Framework Programme for Research and Innovation under the Specific Grant Agreement No. 945539 (Human Brain Project SGA3). In-vivo data was acquired at Scannexus (Maastricht, the Netherlands). OFG is funded by Brain Innovation. We thank Vojta Smekal and Marta Poyo-Solanas for helpful discussions on the involvment of the fronto-parietal network.

## 11 Author Contributions

According to the CRediT system (https://casrai.org/credit/)

**Conceptualization:** A.P., R.G.

**Methodology:** A.P., O.F.G., R.G., I.A.P, G.D.

**Software:** A.P., I.A.P, G.D.

**Validation:** A.P., I.A.P

**Formal Analysis:** A.P.

**Investigation:** A.P.

**Resources:** A.P., O.F.G., R.G., J.P., I.A.P, G.D.

**Data curation:** A.P.

**Writing – original draft:** A.P.

**Writing – review & editing:** A.P., I.A.P, O.F.G., J.P., R.G., G.D.

**Visualization:** A.P., O.F.G., R.G., I.A.P, G.D.

**Supervision:** R.G., J.P., G.D.

**Project administration:** R.G.

**Funding acquisition:** R.G.

## 12 Supplementary Material

**Supplementary Figure 1:**
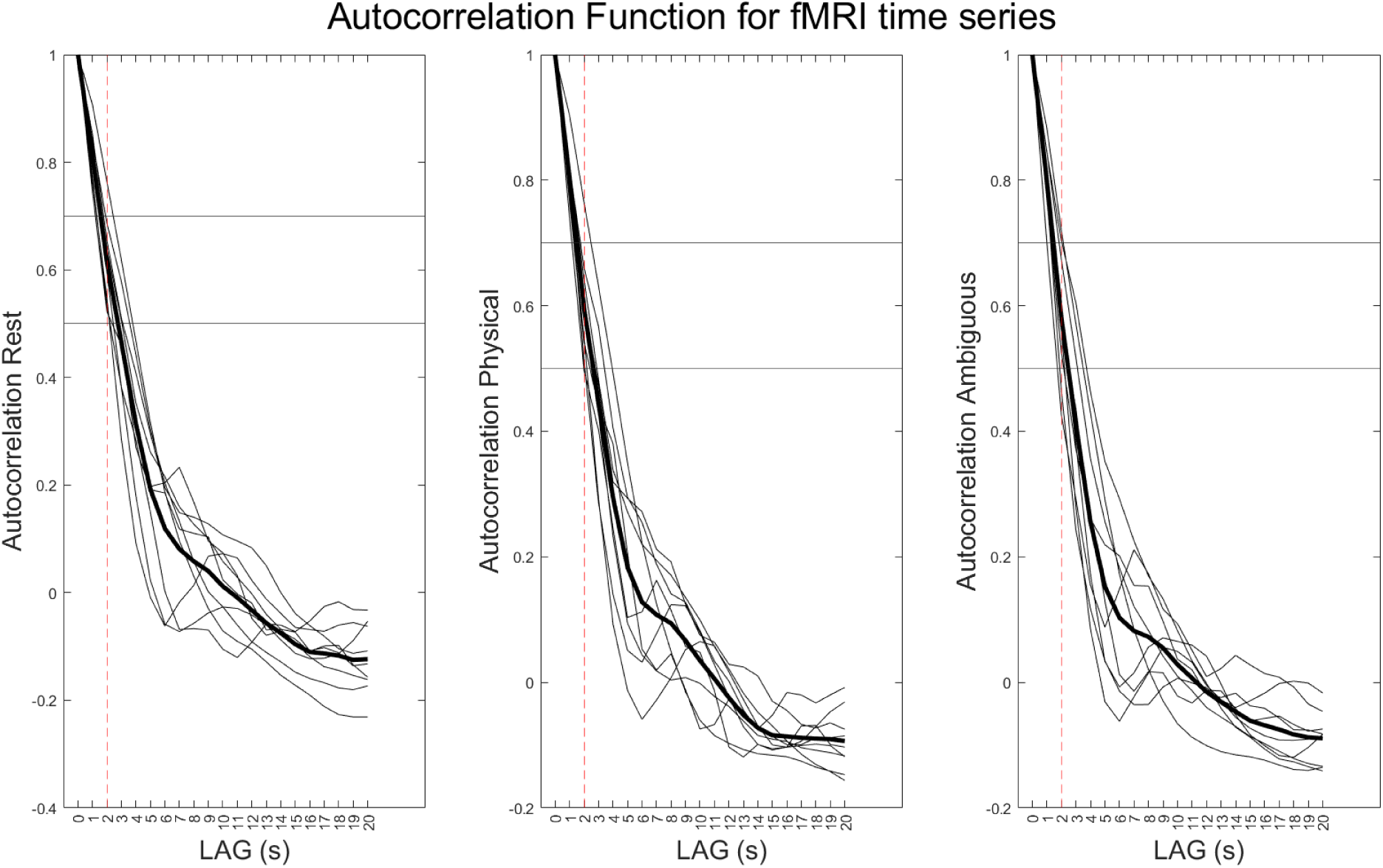
Autocorrelation function (AC) for the resting-state, physical and ambiguous motion condition shown for the first 20 s after condition onset. Thin lines indicate a single-subject AC, while thick lines indicate the average across participants. The AC function consistently decays between 30-50 % (horizontal lines) within our chosen range of lags (1-2-3 seconds).

**Supplementary Figure 2:**
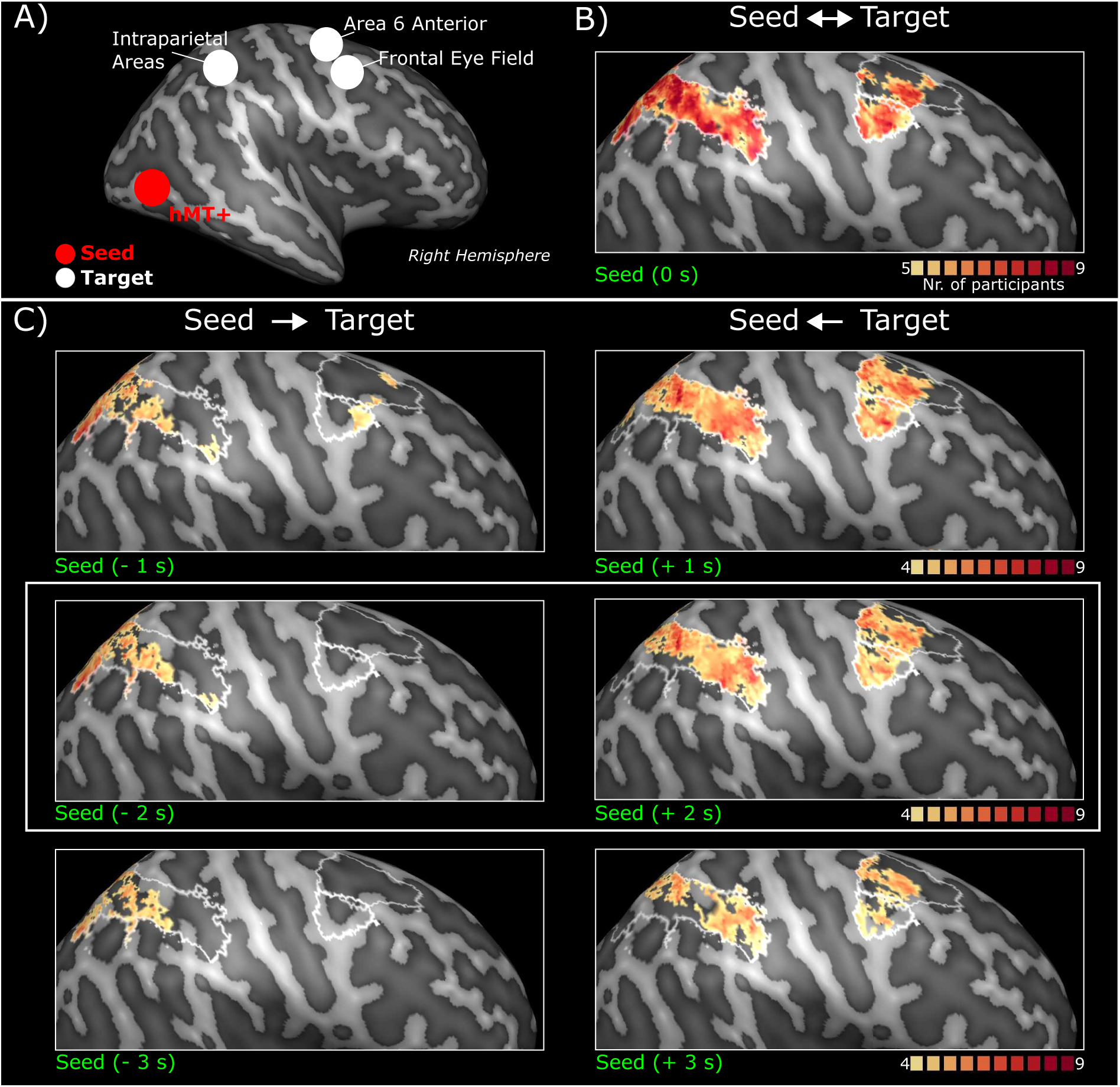
Lag-based functional connectivity analysis for three different lags (+/-1, 2,3 s) computed for the **block-seed hMT+** for the **ambiguous motion** condition. A) Schematic representation of the seed (red voxels within hMT+ indicated by the red circle) and the target areas (indicated by white circles). B) Group conjunction map resulting from correlation analysis (instantaneous synchrony). C) Left column: Seed-to-target group conjunction map. Lag-based FC for backward seed (lag = -2) *>* Lag-based FC for forward seed (lag = +2). Right column: Target-to-seed group conjunction map. Lag-based FC for backward seed (lag = -2) *<* Lag-based FC for forward seed (lag = +2). Mid-row of panel C (white rectangle) corresponds to panel C-D of the main **Figure 5**.

**Supplementary Figure 3:**
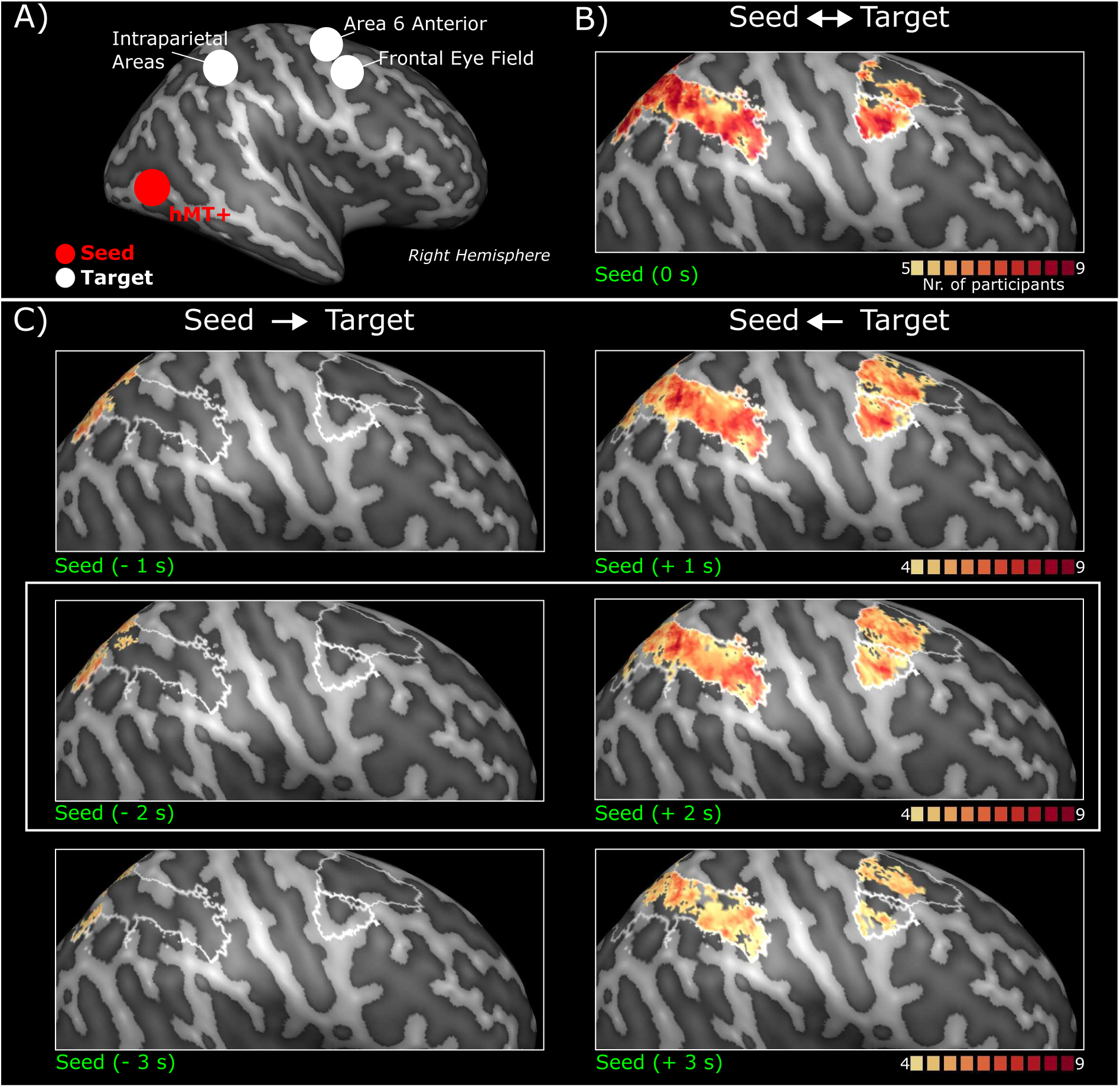
Lag-based functional connectivity analysis for three different lags (+/-1, 2,3 s) computed for the **block-seed hMT+** for the **physical motion** condition. A) Schematic representation of the seed (red voxels within hMT+ indicated by the red circle) and the target areas (indicated by white circles). B) Group conjunction map resulting from correlation analysis (instantaneous synchrony). C) Left column: Seed-to-target group conjunction map. Lag-based FC for backward seed (lag = -2) *>* Lag-based FC for forward seed (lag = +2). Right column: Target-to-seed group conjunction map. Lag-based FC for backward seed (lag = -2) *<* Lag-based FC for forward seed (lag = +2).

**Supplementary Figure 4:**
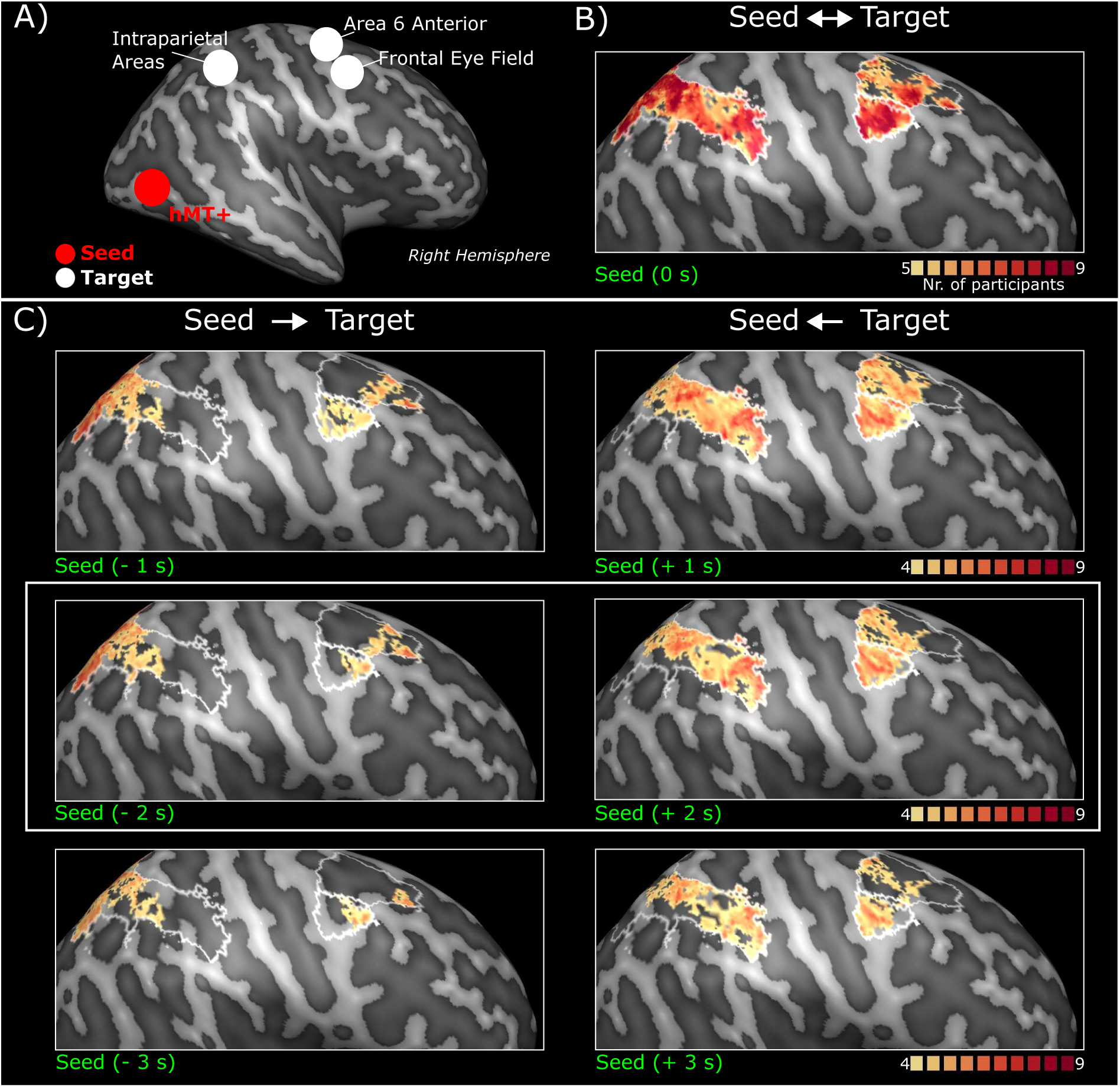
Lag-based functional connectivity analysis for three different lags (+/-1, 2,3 s) computed for the **block-seed hMT+** for the **resting state** condition. A) Schematic representation of the seed (red voxels within hMT+ indicated by the red circle) and the target areas (indicated by white circles). B) Group conjunction map resulting from correlation analysis (instantaneous synchrony). C) Left column: Seed-to-target group conjunction map. Lag-based FC for backward seed (lag = -2) *>* Lag-based FC for forward seed (lag = +2). Right column: Target-to-seed group conjunction map. Lag-based FC for backward seed (lag = -2) *<* Lag-based FC for forward seed (lag = +2).

**Supplementary Figure 5:**
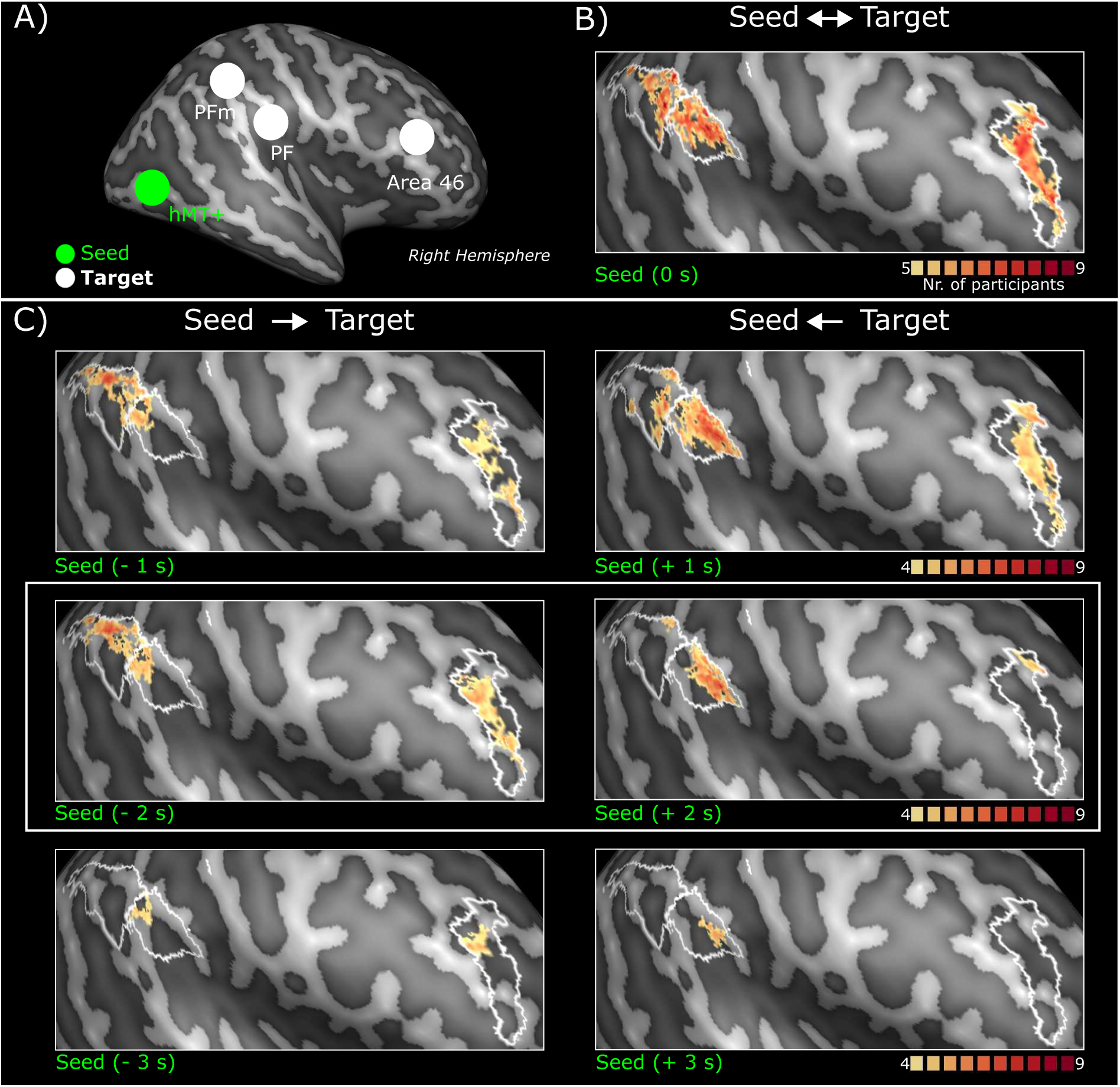
Lag-based functional connectivity analysis for three different lags (+/-1, 2,3 s) computed for the **switch-seed hMT+** for the **ambiguous motion** condition. A) Schematic representation of the seed (green voxels within hMT+ indicated by the green circle) and the target areas (indicated by white circles). B) Group conjunction map resulting from voxel-wise correlation analysis (instantaneous synchrony, i.e. lag = 0). C) Left column: Seed-to-target group conjunction map. Lag-based FC for backward seed (lag = -2) *>* Lag-based FC for forward seed (lag = +2). Right column: Target-to-seed group conjunction map. Lag-based FC for backward seed (lag = -2) *>* Lag-based FC for forward seed (lag = +2). Mid-row of panel C (white rectangle) corresponds to panel C-D of the main **Figure 4**.

**Supplementary Figure 6:**
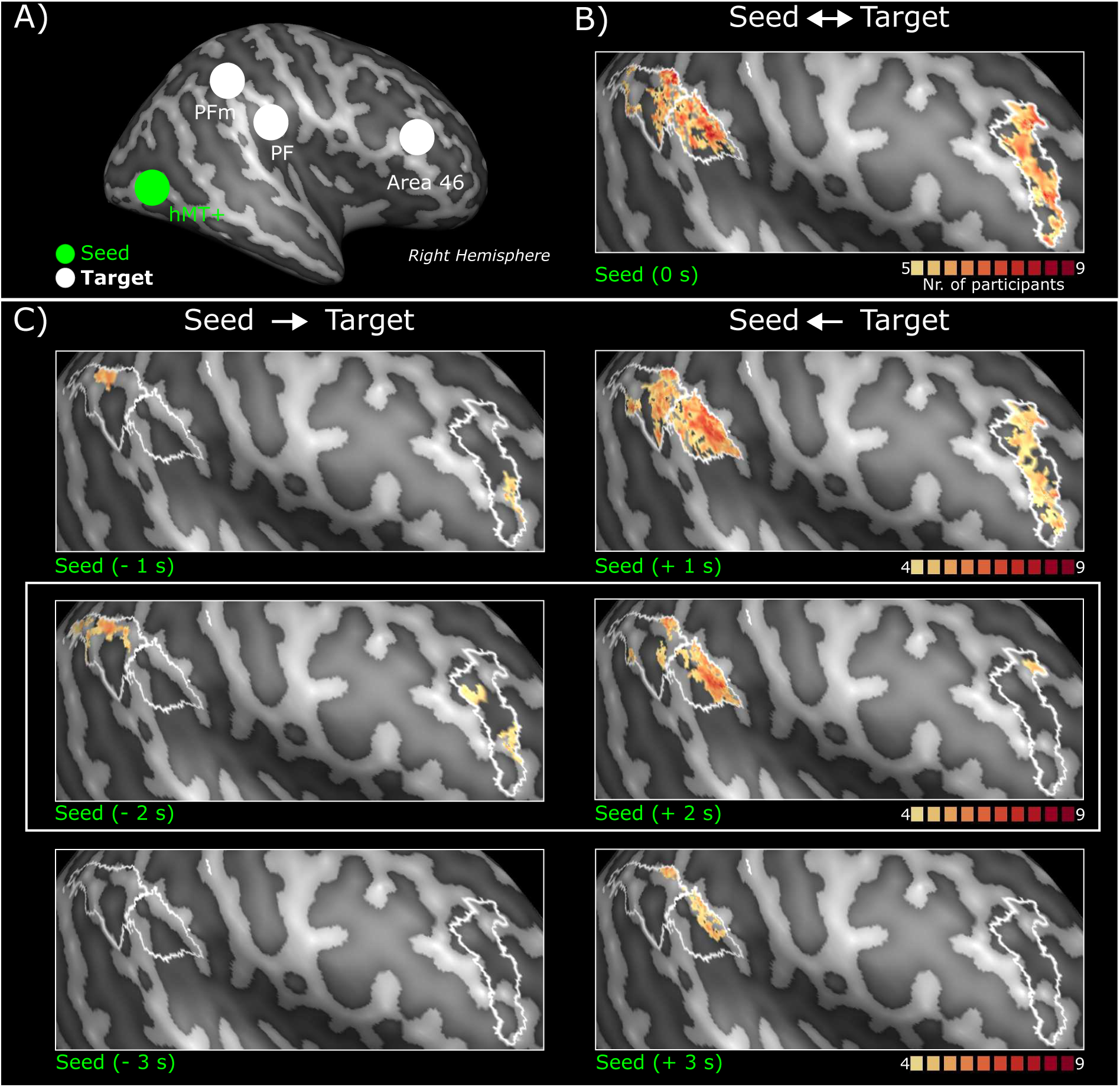
Lag-based functional connectivity analysis for three different lags (+/-1, 2,3 s) computed for the **switch-seed hMT+** for the **physical motion** condition. A) Schematic representation of the seed (green voxels within hMT+ indicated by the green circle) and the target areas (indicated by white circles). B) Group conjunction map resulting from voxel-wise correlation analysis (instantaneous synchrony, i.e. lag = 0). C) Left column: Seed-to-target group conjunction map. Lag-based FC for backward seed (lag = -2) *>* Lag-based FC for forward seed (lag = +2). Right column: Target-to-seed group conjunction map. Lag-based FC for backward seed (lag = -2) *<* Lag-based FC for forward seed (lag = +2).

**Supplementary Figure 7:**
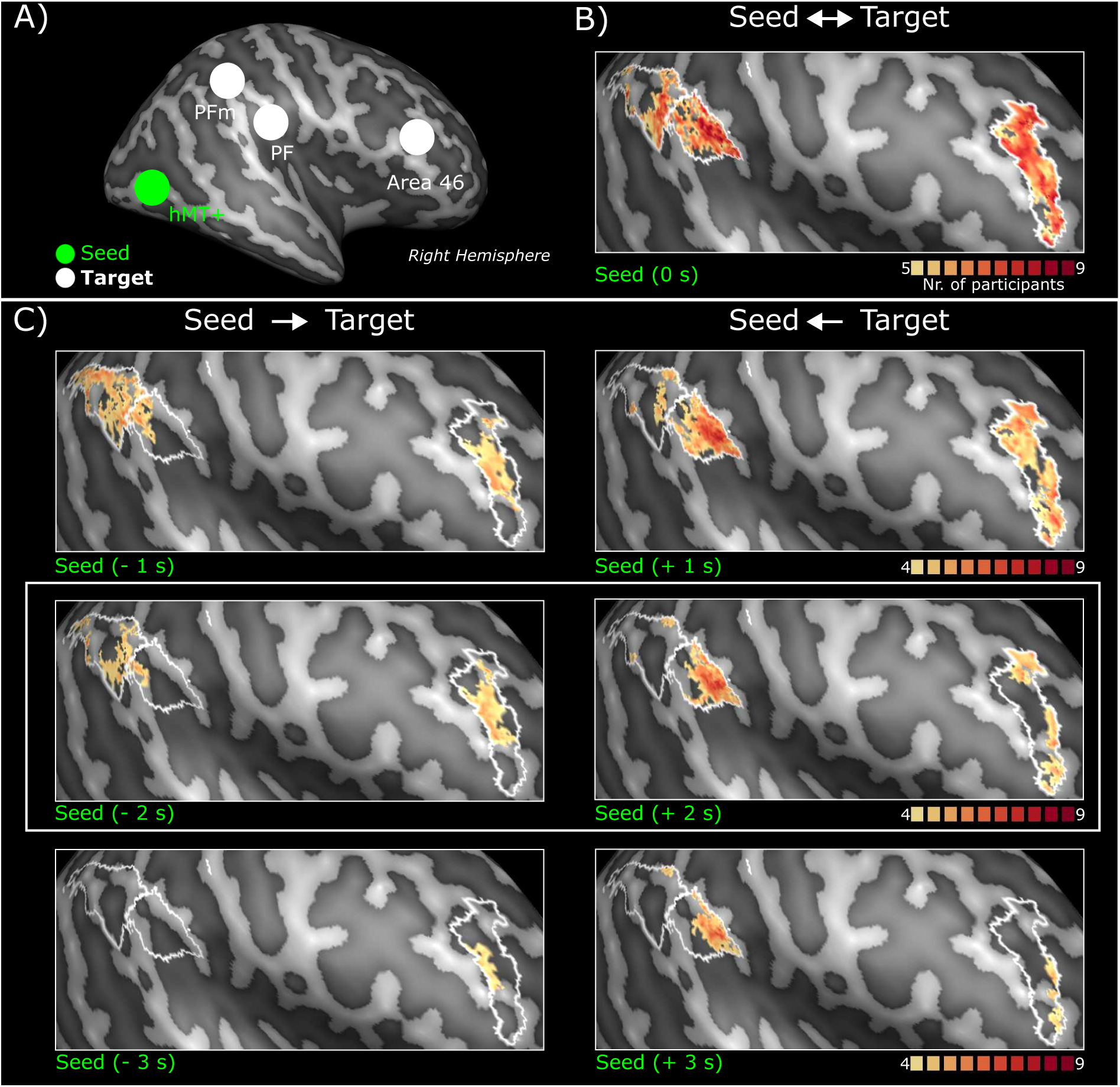
Lag-based functional connectivity analysis for three different lags (+/-1, 2,3 s) computed for the **switch-seed hMT+** for the **resting-state** condition. A) Schematic representation of the seed (green voxels within hMT+ indicated by the green circle) and the target areas (indicated by white circles). B) Group conjunction map resulting from voxel-wise correlation analysis (instantaneous synchrony, i.e. lag = 0). C) Left column: Seed-to-target group conjunction map. Lag-based FC for backward seed (lag = -2) *>* Lag-based FC for forward seed (lag = +2). Right column: Target-to-seed group conjunction map. Lag-based FC for backward seed (lag = -2) *<* Lag-based FC for forward seed (lag = +2).

